# Brain organization of a memory champion

**DOI:** 10.64898/2026.01.11.698919

**Authors:** Roselyne J. Chauvin, Annie Zheng, Athanasia Metoki, Samuel R. Krimmel, Ashley Nielsen, Anxu Wang, Philip N. Cho, Noah J. Baden, Daniel Bower, Kristen M. Scheidter, Julia S. Monk, Forrest I. Whiting, Babatunde Adeyemo, Scott Marek, Benjamin P. Kay, Abraham Z. Snyder, Henry L. Roediger, Kathleen McDermott, Steven M. Nelson, Adrian W. Gilmore, Deanna M. Barch, Timothy O. Laumann, Evan M. Gordon, Nico U.F. Dosenbach

## Abstract

Memory athletes can achieve superior performance (e.g., memorizing 339 digits in 5 minutes) with extensive daily training, by converting abstract information into vivid scenes, and placing them along a mental path, that is later retraced (Method of Loci). Understanding the brain mechanisms underlying such training-derived mastery would increase our understanding of the brain’s memory systems and could suggest novel approaches to improving cognition in other domains. As memory athletes use personalized training techniques, it has been challenging to study them with standard group paradigms. Fortunately, precision functional mapping (PFM) enables detailed investigation of individual brains through repeated sampling of resting-state functional connectivity and task fMRI. Here, we precisely mapped the brain organization of a 6-time U.S. Memory Champion (>13 hours fMRI). Relative to controls, the Memory Champion’s network functional connectivity (FC) was strengthened with the retrosplenial, extrastriate visual, and dorsal frontal cortex (area 55b), as well as with the caudate nucleus. The Memory Champion had modules related to scene and semantic processing not seen in controls, alongside stronger connectivity between the caudate and classical memory networks. During rote memorization, the Champion’s task fMRI patterns were typical, with the hippocampus active during encoding. This pattern was reversed when he used his Method of Loci technique, with greater hippocampal activity during recall than encoding. Hence, intense practice at converting abstract information into more memorable formats can develop a procedural memory skill that utilizes brain regions typically reserved for navigation, language, social cognition, and associative learning.

---

Our ability to memorize is limited^1^ by a small working memory span, historically defined at 7 +/- 2 elements^2,3^. Memory athletes demonstrate that this limitation can be overcome with special techniques and intense training. The 6-time US Memory Champion, Nelson Dellis, for example, was able to memorize the order of a deck of 52 cards in 40.65 seconds, establishing a US record at age 27, in a challenge called ‘speed card’ (Fig. 1a). He is a remarkable example of exceptional adult skill acquisition. While he did not have a great memory when young^4^, as an adult, he became the U.S. Memory Champion within two years of starting training, eventually becoming holder of the most U.S. Memory Champion titles (6x). With continued training, his memorization skills have improved over time. For example, he could memorize 178 digits in 5 minutes at age 27, and 339 digits by age 30 (Fig. 1b).

**Figure 1:**
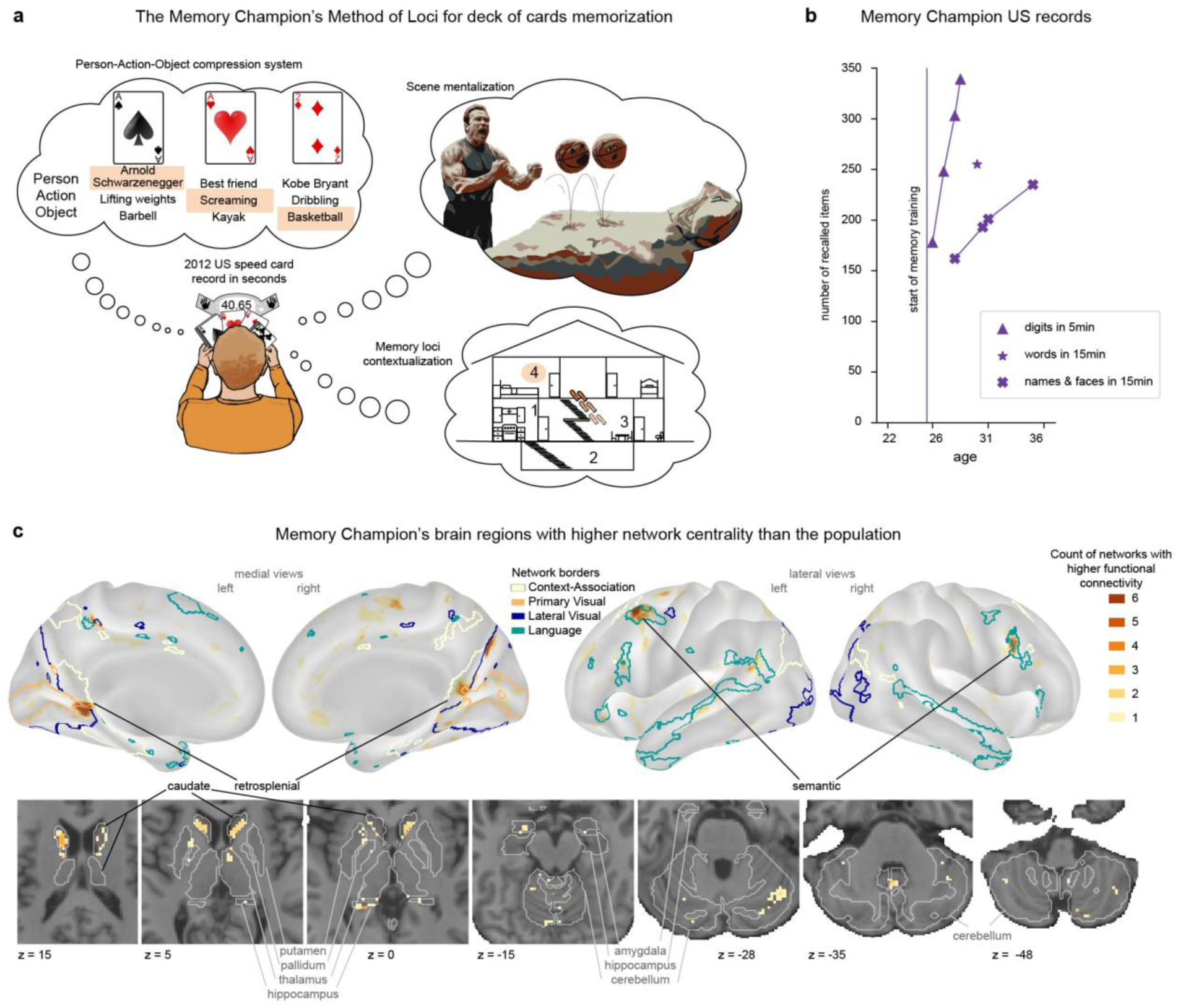
Brain of a Memory Champion: Memorization technique, performance and functional connectivity. **a)** Deck of cards memorization strategy used by the 6x US Memory Champion. The strategy involves a pre-encoded association system where each card is linked to three items (person, action, object), combined into a scene (‘Arnold Schwarzenegger (ace of spades) screaming at (ace of hearts) basketballs (two of diamonds)’) represented at a specific location within one of the Memory Champion’s memory palace loci, e.g. the bedroom of his house. **b)** 6x US Memory Champion’s US records over the years. The number of recalled items for three official US records: digits in 5 minutes, words in 15 minutes, and names and faces in 15 minutes plotted as a function of his age. **c)** Memory Champion’s brain regions with higher network centrality than the population. Regions with higher functional connectivity in the Memory Champion than 95% of individuals from the HCP population (n = 887), at both visits (2015, 2021), with one or more of the 17 functional networks (see Supplementary Fig. 1a) after mixture modeling normalization of each network FC map (see Methods). Darker brown colors indicate stronger FC with a higher number of functional networks (from 1 to 6), indicating higher centrality. Boundaries for the language (teal), lateral visual (dark blue), primary visual (ochre), and context-association (CAN, white) networks are shown in color.

Much of what is known about human memory comes from the study of a single patient (H.M.), who had both hippocampi and surrounding cortex surgically removed to treat his epilepsy^5^. After the surgery H.M. was no longer able to form new declarative memories. However, his procedural memory was relatively preserved, because he steadily improved at repeated tasks (e.g. mirror tracing), even though each time he claimed that it was his first time performing it. Based on H.M. and subsequent work it is known that new information transitions to long-term storage via the hippocampus^6,7^. Several large-scale functional neural networks are important for different aspects of long-term memory and recollection, including the default-mode network (DMN), which is important for self-referential processing^8^, and the parietal memory network (PMN) involved in recall^9,10^. Spatial memory, important for navigation^11^, is supported by the context-association network (CAN) which includes retrosplenial^12,13^, parahippocampal^14^ and entorhinal cortex^15^. Memory consolidation can be enhanced by reward circuitry, including the salience network, which signals the importance of new information^16^ and which was recently found to be closely tied to the PMN^17,18^. Within the reward circuit and basal ganglia, the caudate is particularly involved in linking stimuli and responses during paired association learning^16,19^. The basal ganglia circuit rather than the hippocampus supports non-declarative memory^20^, at least in part explaining the retained procedural learning of patient H.M.

In contrast to individuals with savant syndromes that show remarkable innate memory capabilities^21^, memory athletes use mnemonic techniques and daily training^22^. A core mnemonic technique used by memory athletes is the Method of Loci^23^, in which items to be memorized are placed along a mental path, often referred as ‘memory palace’, that can be mentally navigated to recall information (Fig. 1a). For example, within someone’s house, one could begin in the kitchen, travel to the living room, and then up the stairs to the bedroom, with each location serving as a locus for depositing memories. To increase the number of items memorized (>1 per memory locus), memory athletes use semantic or visual associations and stories to link items. Part of the training is to create an encoding system that is most memorable for them (e.g., types of images, character references). Information can be chunked using a predefined compression system in which each abstract item (such as a card or a digit) is pre-associated with something more easily visualizable (e.g. typically a person, an action, and an object) that are then combined into a visual scene (e.g. person of item 1: ace of spades = Arnold Schwarzenegger, action of item 2: ace of hearts = screaming at, and object of item 3: two of diamonds = basketball) that can be decoded to retrieve information triplets (see Fig. 1a for an example of compression system for cards used by the 6-time US Memory Champion)^24^.

While we understand the techniques that memory athletes use, the brain mechanisms behind their seemingly superhuman performance remain elusive. Prior studies averaged imaging data across groups of memory athletes, an approach that obscures details of functional brain organization^25,26^. Since each memory athlete trains with slightly different techniques, the group level approach is also bound to average across inter-individual training-related differences. Group-averaged studies of memory athletes reported inconsistent observations in retrosplenial cortex with either increased or decreased activity during task performance^27,28^ in addition to reduced fronto-parietal activity^29^ potentially due to practice effects^30^. At rest, group-averaged functional connectivity (FC) differences between memory athletes and controls were reported between memory and visual networks^22^. In the subcortex, FC between the right anterior hippocampus and right caudate has also been found to correlate with memory athletes’ world rankings^31^.

We hypothesized that if a Memory Champion is consistently using his brain in unusual ways, such as cooperative processing between brain regions or networks that do not frequently work together in most people, it would induce Hebbian plasticity^32^. This repeated co-activity between brain regions could lead to increased functional connectivity between them, observable during the resting state, and potentially leading to new functional modules^33^. We therefore tested for the presence of abnormally high cross-network resting-state FC, as indexed by network degree centrality^34^, in the Memory Champion’s brain.

To define individual-specific brain organization, we used a dense repeated sampling approach, called PFM^25,35^ in which large amounts of highest-quality data are acquired for individual-specific analyses^36,37^. PFM has been used to successfully characterize longitudinal brain changes during pregnancy (*n* = 1)^37^, to define cortical and subcortical network organization^15,25,38–40^, to uncover novel networks (i.e., somato-cognitive action network (SCAN))^41,42^, and to determine how they are shaped by plasticity^36,43^. In clinical populations, PFM has been used to identify a promising biomarker of depression^44^, to characterize functional remapping following stroke^45^, and to guide neuromodulation therapies^46^.

For this study, we repeatedly scanned the 6-time US Memory Champion, Nelson Dellis, (in 2021 and 2015) obtaining >400 minutes of resting-state and >350 minutes of task fMRI data. In 2015, he performed the Midnight Scan Club (MSC) protocol^15^, and we used the 10 MSC participants as controls. In 2021, he performed a tailored protocol to map neural circuits underlying his elite memory performance, and two of the MSC participants completed the same protocol to serve as controls (see Table 1 for the list of tasks). We also compared the Memory Champion’s brain, to a large adult population (*n* = 887 participants, 22-35 years old, Human Connectome Project (HCP))^47^ to establish the specificity of his functional brain organization.

**Table 1:** task and rest numbers of runs per participant.

| task | MC | MSC02 | MSC06 |
| --- | --- | --- | --- |
| Rest (15 minutes) | 7 | 5 | 4 |
| Deck of cards memory task (15min) | 8 | - | - |
| Pi retrieval (15min) | 5 | - | - |
| Reading span (10min) | 7 | 9 | 8 |
| Stanford VPN Localizer (8 min) | 7 | - | - |
| Action localizer (7 min) | 8 | - | - |

## Functional connectivity specific to the Memory Champion

Following the hypothesis that intensive Method of Loci training might strengthen cross-network FC via Hebbian plasticity, we compared the Memory Champions FC maps to those of 887 controls (HCP: aged 22-35). For this comparison, we used 17 canonical functional networks (shown in Supplementary Fig. 1a; see specificity analysis in Methods).

For each network we identified brain regions where the Memory Champion’s functional connectivity was stronger than 95% of the controls. Several brain regions exhibited greater FC compared to controls with six or more functional networks at both timepoints (2015 and 2021 visits), elevating the centrality of these regions and thus making them candidate regions for memory training effects (Fig. 1c; Table E1).

Cortical regions with the highest centrality in the Memory Champion relative to controls included retrosplenial (context-association network), parieto-occipital (higher-order lateral visual), and frontal regions belonging to the language network, particularly those previously associated with semantic processing^48^ (including Brodmann area 55b). In subcortex, regions with the greatest centrality increases relative to controls were anterior caudate and central cerebellum. The caudate showed stronger FC with memory and executive networks (PMN, DMN, CAN, fronto-parietal network (FPN), Salience). In contrast, all cortical and the cerebellar high centrality regions showed greater FC mainly with visual and action control networks (Table E1).

The same analyses performed on two matched controls (MSC02, MSC06), failed to reveal strong divergence from the HCP group data, in that no brain region showed increased RSFC with more than 3 networks. Moreover, the regions with the relatively greatest differences from the HCP group data in the controls (MSC02, 06) did not overlap with the increased centrality regions in the Memory Champion (Supplementary Fig. 2).

## High centrality regions connected to vision, action and memory

To visualize the details of functional connectivity differences between the Memory Champion and matched repeatedly sampled controls (Fig. 2 for the 2021 visit, see Extended Data Fig. 2 for the 2015 data and Supplementary Fig. 3 for additional brain views), we computed correlation seed maps for regions with the greatest centrality increases in the Memory Champion. We selected matching regions in the Memory Champion and control participants (MSC02, MSC06): retrosplenial cortex (Brodmann areas 26, 29, and 30^13,49^), a left semantic region (Brodmann area 55b based on individual language network functional segmentation, see Extended Data Fig. 1, see Methods), and the head of the caudate (from individual anatomical segmentation, see Methods). Individual-specific seed maps were generated for each of these regions for the Memory Champion (Fig. 2, left) and the controls (Fig. 2, middle & right; see Extended Data Fig. 2 for quantification of individual resting state scan distribution and t-test between Memory Athlete and PFM controls in both 2015 and 2021 datasets).

**Figure 2:**
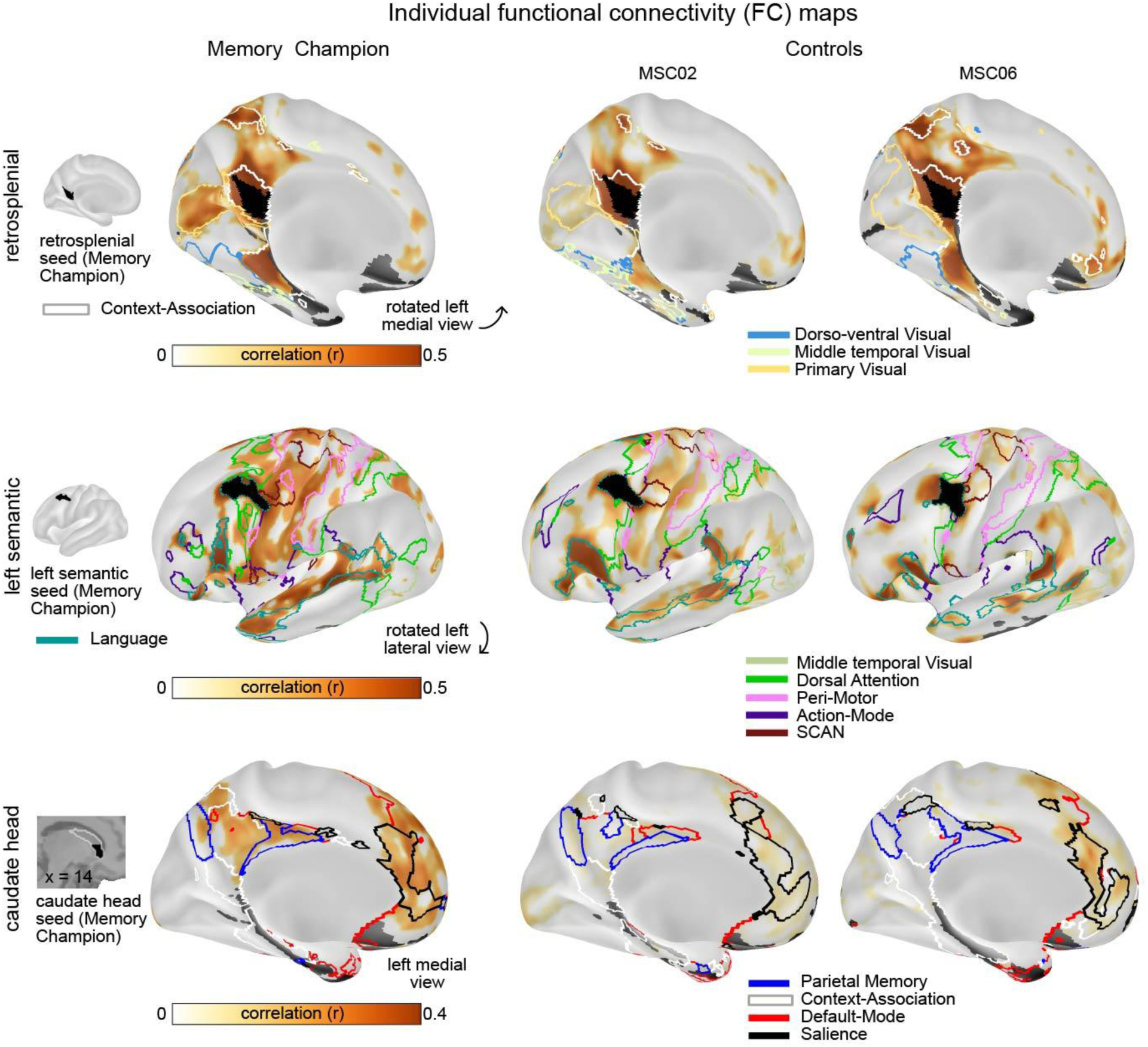
Functional connectivity of regions with high centrality in the Memory Champion. Whole-brain FC maps for representative brain regions identified in Fig. 1c, top 20% of strongest connections are shown. Left column shows the Memory Champion, middle column MSC02 control, and right column MSC06 control (2021 data; see Extended Data Fig. 2 for 2015 data, and Supplementary Fig. 3 for additional brain views). The top row shows correlation (r) for retrosplenial cortex (Brodmann areas 26, 29, and 30, part of context-association network -white border), middle row shows the left semantic region (area 55b; language network -teal border) and the bottom row shows the head of the caudate. The seed regions are shown in black. All other network borders indicate networks significantly more connected in the Memory Champion compared to the controls in 2021 (see Extended Data Fig. 2 for quantification). SCAN stands for somato-cognitive action network. Low signal-to-noise masked areas are displayed in dark grey.

The retrosplenial region showed significantly greater FC with visual networks in the Memory Champion compared to controls, particularly with the primary visual and dorso-ventral stream networks (Fig. 2; Extended Data Fig. 2 top rows, one-tailed independent *t > 4.2, P <* 0.001, FDR-corrected). The left semantic region had significantly greater FC than controls with a set of networks involved in motion and action (Fig. 2; Extended Data Fig. 2 middle rows, one-tailed independent *t >* 2.6, *P <* 0.03, FDR-corrected), namely the visual medial temporal network, which includes visual area 5 associated with motion perception^50^, the dorsal attention network (active in the Memory Champion during action imagery, Supplementary Fig. 4), and the peri-motor, action-mode, and somato-cognitive action networks^51^. In the Memory Champion, the caudate head showed significantly stronger FC with memory networks (parietal memory and context association networks) and the salience network (Fig. 2; Extended Data Fig. 2 bottom rows, one-tailed independent *t > 2.4, P <* 0.03, FDR-corrected). Stronger FC with the default-mode network was only significant in the 2021 dataset (*t = 17.8, P <* 10^-9^, FDR-corrected). We also validated that similar results were observed when testing exploratively across all networks (Extended Data Fig. 2c).

## Scene and semantic modules of the Memory Champion

To further characterize the high centrality regions (Fig. 3a), we annotated them based on their network labels and then grouped them into modules based on functional similarities. One module belonged to the visual and context association networks, including the retrosplenial cortex, parietal occipital sulcus, middle and inferior temporal regions, and posterior intraparietal sulcus. The other module was part of language, auditory, executive (fronto-parietal and dorsal attention), and motor (somato-cognitive action and motor hand) networks, which included frontal regions from the semantic part of the language network, anterior precuneus, paracentral lobule, superior temporal regions, and anterior intraparietal sulcus (see Table E1 for coordinates).

**Figure 3:**
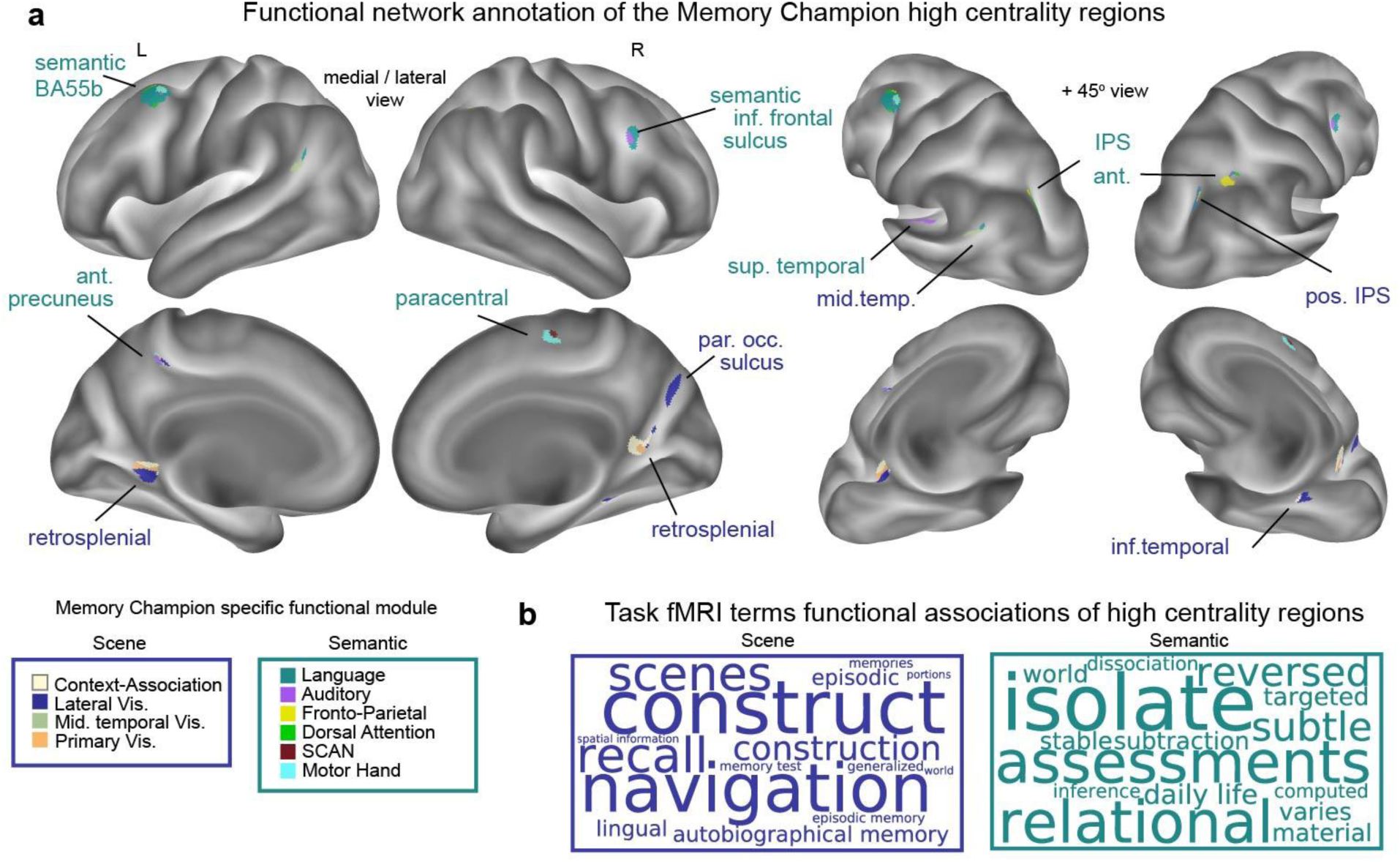
Functional modules only found in the Memory Champion. **a)** Functional network annotation of the Memory Champion’s high centrality regions. The high centrality regions from Fig. 1c, are shown on an inflated brain, colored by their functional network assignments. Regions are grouped into two sets of networks (network color label, bottom left) representing a scene (dark blue) and semantic (teal) module. Each region is named anatomically or functionally in dark blue for the scene module and teal for the semantic module. Abbreviations: parietal occipital (par. occ.), intra parietal sulcus (IPS), somato-cognitive action network (SCAN), Brodmann Area (BA). **b)** Top 15 meta-analytic task fMRI database terms (Neurosynth^52^) associated with the Memory Champion’s high centrality regions. The font size of the word in the word clouds matches the relative weight of the term amongst the top 15.

To investigate what is known about the high centrality regions of the Memory Champion, we annotated them using the automated meta-analytic Neurosynth term maps^52^, which are based on activation peak coordinates from a large corpus of task fMRI studies (Fig. 3b for module-level analysis and Extended Data Fig. 3 for replication at the individual region level). The high centrality regions contained in visual and context association networks were associated with terms related to scene representation: construct, navigation, scene, and recall. Therefore, we named this set of regions the ‘scene module’. The other set of high centrality regions was associated with terms related to abstract concept manipulation: isolate, assessment, relational, and reversed. Since these terms refer to abstract, analytic, or experiential concepts that are generally higher-level rather than sensory or formal, and several of these regions were within the semantic part of the language network, we named this set of regions the ‘semantic module.’

The scene and semantic modules specific to the Memory Champion also emerged from the data when using other analytic approaches, such as principal component analysis on high centrality region FC maps, testing the functional similarity of high centrality regions against null hypotheses, and exploring module differences in network connectivity (see Module Analysis, Methods; Extended Data Fig. 4).

## Caudate more strongly connected to memory networks

To further analyze the FC of the Memory Champion’s high centrality regions to subcortex, we mapped out voxels where his FC was stronger than 95% of the controls (HCP), for each of the memory networks (DMN, PMN, CAN) (Fig. 4a; for replication in 2015 data see Supplementary Fig. 5). The Memory Champion had stronger FC between the striatum and all memory networks, particularly in the caudate (FC > 100% of controls). The head of the Memory Champion’s caudate was more strongly connected to all three memory networks (Fig. 4a; replication with 2015 data in Extended Data Fig. 5).

**Figure 4:**
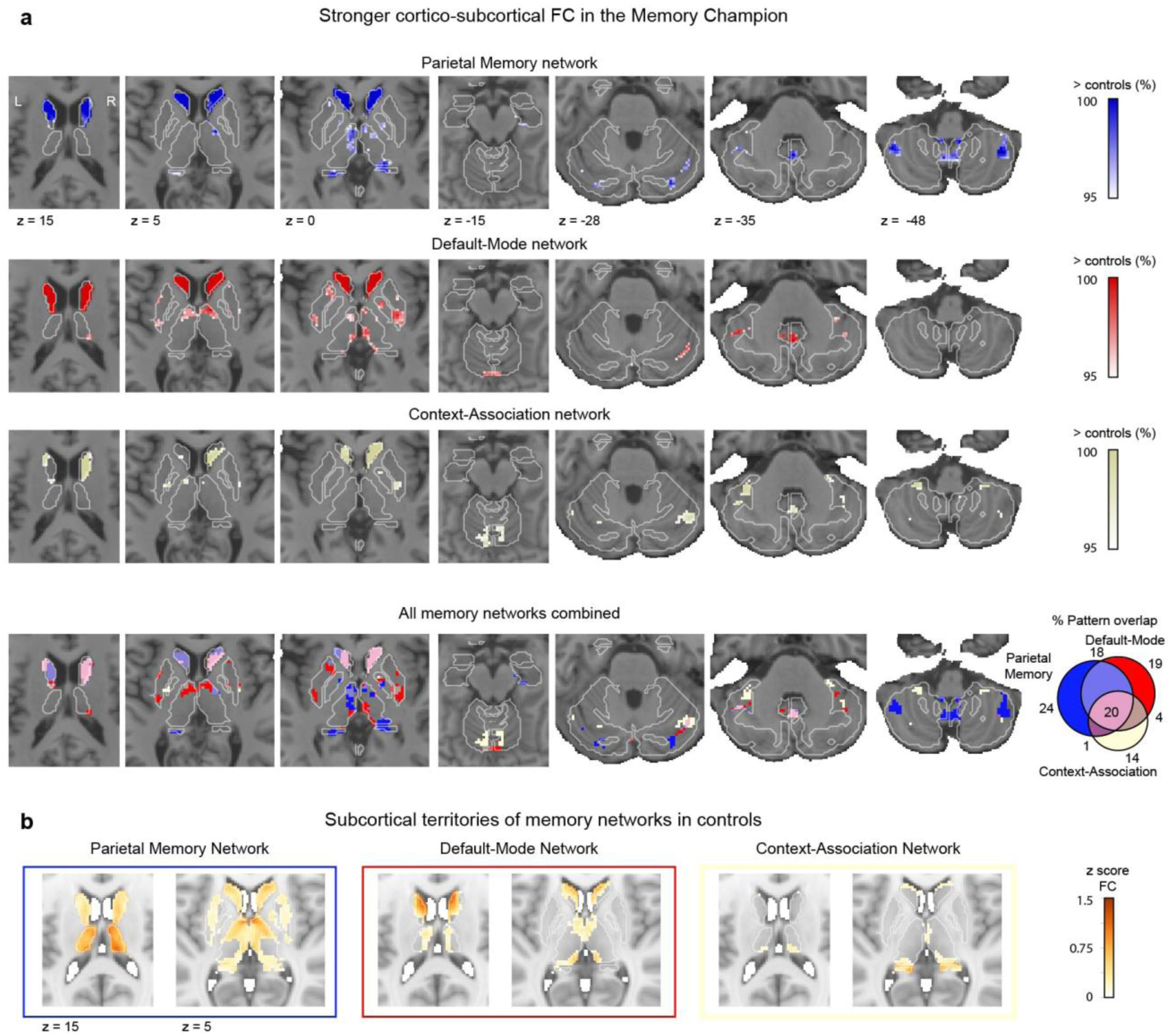
Memory networks to subcortex functional connectivity. **a)** Subcortical regions where the Memory Champion showed stronger FC than 95% of controls (HCP, n = 887), by network: top row parietal memory network (blue), second row default-mode network (red) and third row context-association network (white) FC maps in subcortex. The overlap across networks is shown in the fourth row, with pink indicating greater functional connectivity to all three memory networks. Results are displayed on the Memory Champion’s horizontal T1 structural slices with the network labelled color as gradient scale (right). White borders outline the Freesurfer segmented anatomical structures. See Extended Data Fig. 5; Supplementary Fig. 5 for replication in 2015 data. **b)** Subcortical functional connectivity of memory networks in controls (HCP). Left to right, row parietal memory network (blue), default-mode network (red) and context-association network (white) Z scored FC maps for regions where networks are competing for winner-take-all labels (i.e. network territories). Results are displayed on the MNI horizontal T1 structural slices. White borders outline anatomical structures. (See Extended Data Fig. 6 for winner-take-all maps, Extended Data Fig. 7 for additional slices of network territories in controls and quantification of pattern overlap with the stronger cortico-subcortical FC area in the Memory Champion. See Supplementary Fig. 6 for subcortical network territories of the Memory Champion).

In addition to generally stronger connectivity to the striatum, we found that the Memory Champion had a pattern of subcortical functional connectivity in relation to memory systems that was distinct from controls. Controls did not exhibit FC between caudate and the context-association network contrary to the Memory Champion (See Fig. 4a third raw and Fig. 4b third column). Generally, the Memory Champion showed stronger FC between subcortex and memory networks in subcortical regions connected to fronto-parietal, parietal memory and salience networks in controls (Extended Data Fig. 7c, one-tailed testing against null, *P <* 0.05, FDR-corrected). This suggests that the Memory Champion has not only strengthened existing functional connections (parietal memory network) but also possesses connectivity absent in controls, such as that between the caudate and context-association network.

## Contrasting activity patterns during rote memorization and Method of Loci

To understand whether the Memory Champion’s high centrality regions are related to his memory skills, we analyzed task fMRI activity during the performance of two memory tasks. One task was a working memory task (reading span) performed by controls and the Memory Champion using rote memorization; the Memory Champion did not use any memory techniques. Participants engaged in rote rehearsal of the items (3 to 7 words, encoding) before being cued to silently recall them (proactive recall) (Fig. 5a).

**Figure 5:**
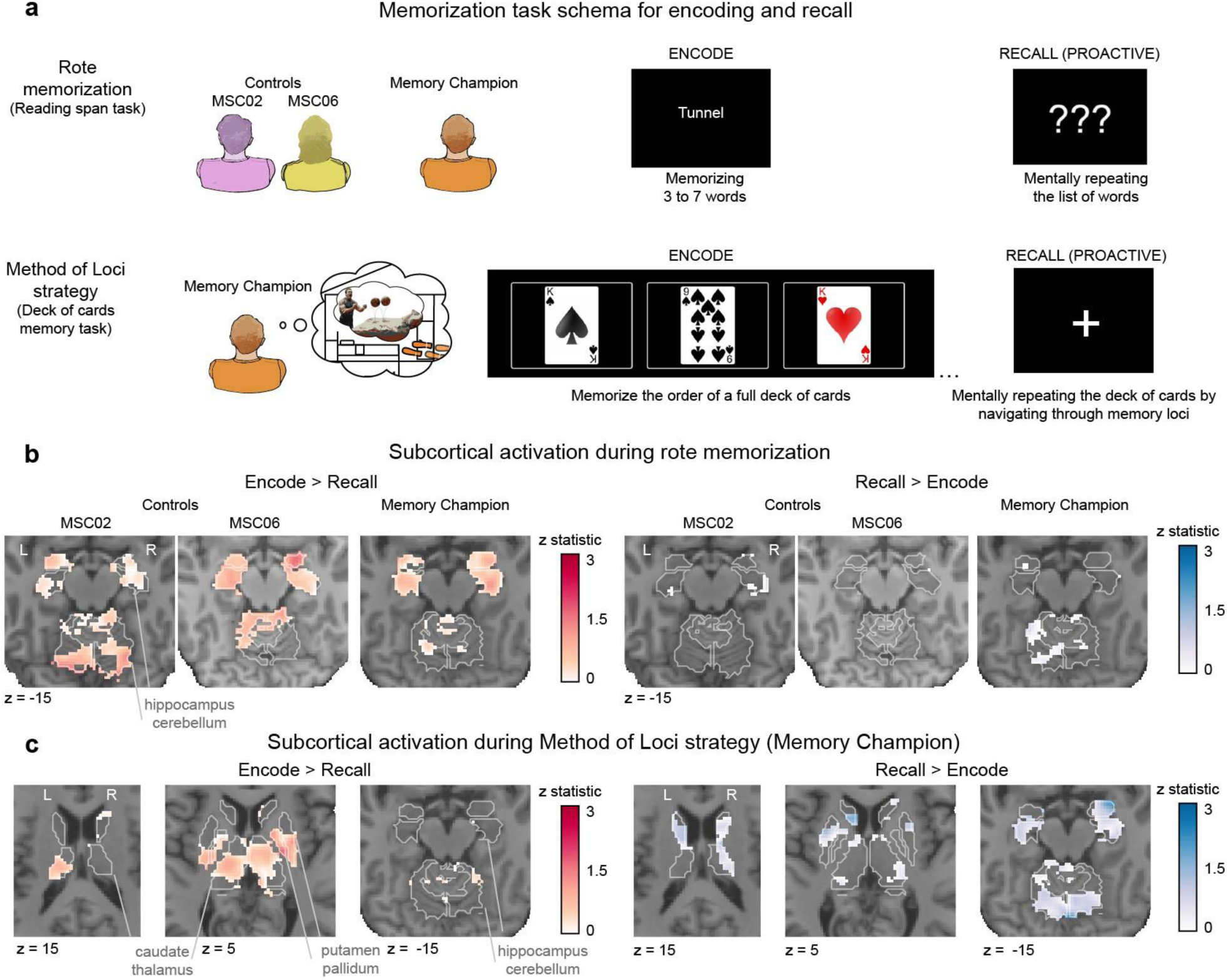
Task activations during encoding and recall when using rote memorization or Method of Loci. **a)** Reading span and Deck of cards memory task schemas for encoding and recall (proactive). Schema shows the encoding and proactive recall phases of the reading span task (top line) and the Deck of cards memory task (bottom line). Full task schemes are available in Supplementary Figs. 7 and 8. **b)** Task fMRI contrast of Encoding versus Recall, when using rote memorization in the Reading span task. Results are shown on each participant’s T1 horizontal slice (z = −15) for unthresholded Z-statistic maps of Encode > Recall (left) and Recall > Encode (right), for controls (MSC02, MSC06) and the Memory Champion (2021 data). White borders outline anatomical structures (Freesurfer segmentation). For additional slices and cortical results see Extended Data Fig. 8. **c)** Task fMRI activation of Encode versus Recall contrasts when the Memory Champion is using the Method of Loci strategy in the Deck of cards task. Controls are unable to perform this task. Results are shown on the Memory Champion’s T1 horizontal slices (z = 15, 5, −15) for unthresholded Z-statistic maps of Encode > Recall (left) and Recall > Encode (right), with white borders of anatomical structures. For cortical results see Extended Data Fig. 9.

The second task was a custom-tailored task that only the Memory Champion could perform, in which he used the Method of Loci and his Person-Action-Object compression system (Fig. 1a) to memorize the order of a deck of cards (Deck of cards memory task, Method; Fig. 5a). The encoding phase, during which the deck was displayed 3 cards at a time for 6 seconds, was followed by a proactive recall phase, in which the Memory Champion retrieved the order of the deck of cards just memorized by navigating his memory loci. Correctness of the memorized deck order was verified outside the scanner. We compared encoding and recall neural activity in both tasks to capture the neural circuit underlying his superior memory performance.

During rote memorization (Fig. 5b) controls (MSC02 and MSC06) and the Memory Champion showed similar activations in subcortex (Fig. 5b) and cortex (Extended Data Fig. 8). This pattern of activation was consistent with previously published work^53,54^, including stronger activation of the DMN during encoding and PMN during recall. Rote memorization elicited more hippocampal activity during encoding than proactive recall, again consistent with prior work^55^.

In sharp contrast, when using the Method of Loci, the Memory Champion showed the reverse pattern, with greater hippocampal activity during recall than encoding (Fig. 5c) and deactivation during encoding (Extended Data Fig. 9a). Cortical activations were also very different between the two strategies (Extended Data Fig. 9b; Supplementary Fig. 9), including frontal activity in Method of Loci consistent with strategy based memorization^56^. Both encoding and proactive recall elicited activity in the caudate during Method of Loci memorization, highlighting the importance of the caudate in this mnemonic strategy.

To evaluate whether the Method of Loci method utilized the high centrality regions identified by FC, we quantified their mean activation in encoding versus recall contrasts for both rote memorization and the Method of Loci task (Supplementary Fig. 10) and noted greater activation overall for the Method of Loci (Encoding (E)>Recall(R) mean (sd) *t* = 0.72 (1.92), with significant right posterior IPS activity, *t* = 7.98, *P* < 0.001, one-tailed independent t-test, FDR-corrected; R>E *t* = 0.48 (0.51)) compared to rote memorization (E>R mean *t* = 0.09 (0.15), R>E mean *t* = 0.08 (0.09)).

## Regions active during Method of Loci training show altered FC at rest

To better understand whether the Method of Loci training may have contributed to the exceptionally high centrality of some regions, we designed a task based on a memory challenge that the Memory Champion was preparing for at the time: fast retrieval of the locations of 5-digit sequences within the first 10,000 digits of pi. Having already memorized the first 10,000 digits of pi, he was presented with a series of 5-digit sequences and asked to retrieve their locations within the overall sequence of pi (Fig. 6a). Testing of recall location of each sequence was verified outside of the scanner. Failed and successful retrievals within 6 seconds were evenly represented as he recently had retrained the first 5000 digits. We contrasted the neural activation during failed trials (search only: ‘Search’) and successful trials (search + found: ‘Found’). The search process significantly activated the dorsal attention network and an extended portion of the CAN (Fig. 6b and c, test against null distribution, *P* < 0.05, FDR-corrected). Finding the location of the 5-digit sequence was associated with greater activation of the visual cortex and showed significant activation of several high centrality regions, including the parietal occipital sulcus and the intraparietal sulcus (IPS) regions (Fig. 6d and e, test against null distribution, *P* < 0.05, FDR-corrected). These results indicate that the Memory Champion’s practice methods engage some of the same brain regions that exhibited altered functional connectivity (higher centrality relative to controls) in the resting state.

**Figure 6:**
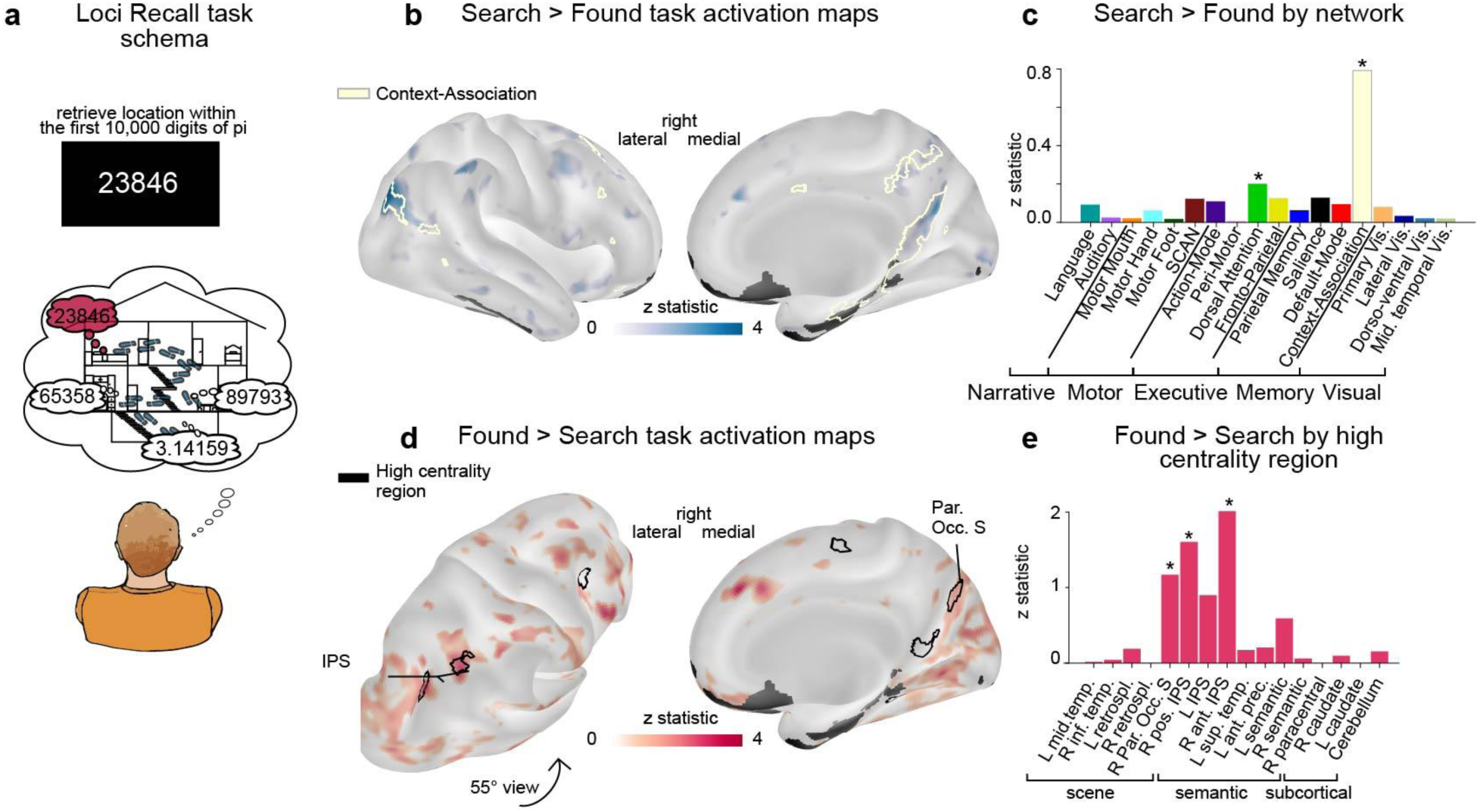
Task fMRI activation when recalling digits of the number pi using Method of Loci. **a)** Task schema for recalling the location of digit sequences within the number pi. Mental navigation (search; in blue) and retrieval (found; in red) of the locus where a prompted unique 5-digits sequence from the pre-learned 10,000 first digits of the number pi is stored. Contrasted events are ‘search’ (failed to find within 6 seconds) and ‘found’ (localized within 6 seconds) events. (see Supplementary Fig. 11 for full task schema). **b)** Memory Champion task activation contrast of Search > Found recall of 5-digit sequence locations within the 10,000 first digits of the number pi. Context-association network borders (white outlines) are displayed. **c)** Search > Found activations quantified by functional network. Average Z-statistic bar plot (*, P < 0.05, FDR-corrected). **d)** Memory Champion task activation contrast of Found > Search recall of 5-digit sequence locations within the 10,000 first digits of the number pi. Memory Champion high centrality region borders (black outlines) are displayed. **e)** Average Z statistics per high centrality regions grouped by scene and semantic modules and subcortical regions. Low signal-to-noise masked areas are displayed in grey. See additional views, network and region bar plots in Extended data Fig.10

## Accessing domain-specific memory circuits

To an external observer, the Method of Loci is indistinguishable from performing rote memorization. Yet, precision brain imaging revealed that Method of Loci strategies are supported by a fundamentally different set of cortical and subcortical regions than rote memorization (Fig. 5; Extended Data Fig. 8), forming new circuits that can be detected at rest.

During rote memorization, hippocampal activity is so tightly linked to encoding that it predicts recall success^57–59^. When using rote memorization, the Memory Champion showed the same activity patterns as controls, relying on classical memory networks (default-mode^8^, parietal memory^9,10^), for encoding and recall (Extended Data Fig. 8; Supplementary Fig. 10), with the hippocampus most active during encoding. When using the Method of Loci, the Memory Champion’s hippocampus was surprisingly inactive, which could indicate that little effort is required on his part despite the nominal complexity of the strategy. Alternatively, Method of Loci encoding might depend more on the caudate than the hippocampus. Afterall, H.M. still had procedural memory and could learn new skills without his bilateral medial temporal lobes^5^. Under either scenario, the findings indicate that Nelson Dellis did not supercharge his hippocampus-based memory circuits or improve his general-purpose memory to become a Memory Champion. Instead, his technique, which converts abstract information into more naturally memorable formats, allows him to access the brain’s much larger domain-specific memory for landmark-based navigation and storytelling.

It is perhaps unsurprising that human brains can solve a basic problem, such as remembering sequences, in multiple ways. Yet it is striking that switching from a general-purpose system that can hold information of any type to those specialized for navigation and social communication, improves speed and capacity so massively (e.g. 339 digits in 5 minutes; 10,000 digits of pi). What appears like improved working memory depends on conversion strategies that containerize abstract information in such a way that circuits for procedural learning, as well as spatial and social cognition can be accessed. These results suggest that while our ability to rote memorize abstract information maxes out at a phone number, we have the capacity to encode a thousand times more information if it helps us find our way (navigation) or build relationships (social, communicative).

## Caudate for memory as a skill

The caudate helps strengthen correct associations^60^ by producing^61^ and evaluating^62^ confidence values during learning^63^. The provision of such feedback signals during associative learning is thought to make the caudate critically important for skill acquisition, and habit formation^19^. The Method of Loci builds on multiple paired association strategies. Some paired associations are pre-encoded if-then structures: a sequence of environmental cues and motor actions to navigate the memory palace (“if kitchen, turns right to the leaving room”, using egocentric navigation^64^) and a compression-system (if person, action, object then abstract information (card/digit)), while some paired associations are built for the specific memorization task (if loci then scene). The path through the memory palace is used as a model to rebuild associations, with the caudate being active during both the encoding and recall phases of Method of Loci performance (Fig. 5c). The strengthened connectivity between the caudate and memory networks underlines its key role in the Method of Loci. Memory athletes are tested on the quantity and speed of memorization. We hypothesize that the caudate might play an important role in speeding up memorization by making the rapid formation of new associations (e.g., between a locus in a memory palace and a triplet of cards) a skill^11,60^. The effectiveness of repeated testing compared to repeated studying^65^ suggests that paired-association and feedback are generally advantageous to long term learning.

Declarative (medial temporal lobe system) and non-declarative (basal ganglia system) memory processes interact or compete depending on the learning task^20^. During skill learning, they interplay, starting with more declarative processing^66^ and moving toward automatization^67^ and skill consolidation^68^, which is sensitive to declarative interference^69^. The strengthened connectivity between memory networks and the caudate implying a consolidated skill, consistent with reduced hippocampal activity during Method of Loci encoding (Fig. 5c). As the Memory Champion wrote: *“After years of using these techniques, my [mental] images boil down to a nuance that I can’t explain in writing. It’s almost like a feeling”*^24^, suggesting that he has created a cognitive containerization skill that encodes abstract information in a non-declarative format, which he decodes at recall. Thus, highlighting the advantage of non-declarative memory.

## Context and narration enhance memory span

During the Method of Loci visual and context-association network activity was greater during encoding than recall (Extended Data Fig. 9), consistent with Nelson Dellis’ strategy based on scene building and visualization with person-action-object triplets deposited in specific memory palace loci. The retrosplenial cortex, important for searching for information in a memory palace (Fig. 6), was also active when the Memory Champion looked at physical places (Supplementary Fig. 4), consistent with its known role in spatial contextualization^12,70^. The broad activation of visual regions and not typical memory networks (default-mode, parietal memory) coincides with the Memory Champion’s reported use of concrete imagery in his rapid encoding, suggesting that he uses the same brain regions for memorizing cards and digits, as he does to encode and remember true visual scenes (Extended Data Fig. 9; Fig. 6c).

The brain regions constituting the semantic module showed increased FC with action networks (Table E1). Particularly the left semantic region, which includes Brodmann area 55b, is one of the language regions that develop later in children^71^, known to be active in story listening^72^, specifically processing information related to syntax and story event boundaries^73^. We hypothesized that the semantic module is involved in tracking the story elements across memory loci, particularly the action aspects. As the Memory Champion explains, *“a true, powerful, memorable mental image needs to be created with all of the senses, not just mental sight. […] Lastly, […] add movement…. For whatever reason, things that move are difficult to forget. My theory is that by making it move, you’ve suddenly made that still photo in your mind an actual story”*^4^. Indeed, actions (memory for actions, enacted events) play a critical role in memory processes by providing rich, multifaceted encoding experiences that enhance the durability and retrieval of memories^74^. Prior to the invention of writing systems, complex histories such as Homer’s Odyssey^75^, would be transmitted as stories and songs. Evidence of large knowledge memorization relating ancient historical events was also demonstrated in the oral culture of Australian Aboriginal communities^76^, suggesting deliberate use of narration for its superior memorizability^77^. Information is more easily recalled when embedded in a coherent and attention-grabbing narrative, even when told only once. For example, nobody fears that their credit card number will be remembered by someone having heard it only once. In sharp contrast we fully expect someone to remember a 16-byte story about an escalating conflict between two coworkers. Hence, a key component of the superior abilities of memory athletes may be their capacity to convert arbitrary information into easily imageable scenes^78^ that help create a more memorable narrative structure^79^.

## Module based enhanced cognition

It remains possible that Nelson Dellis and other memory athletes were born with an innate talent for memorization using the Method of Loci. However, the improvement in memory performance over the years (Fig. 1b), and the connectivity of the semantic and scene modules (Fig. 6c), suggest that the specific patterns of connectivity in the Memory Champion’s brain were more likely shaped by repeated co-activation during memory training. Together with the evidence of late plasticity of Brodmann area 55b in development^71^, these regions may flexibly combine into processing modules that support strategies which enhance cognition. Much like neural plasticity in brain regions for reading^80^ and writing^81^, specific mnemonic strategies like the Method of Loci may significantly improve learning and memory capabilities by turning a set of regions into a new module supporting a cognitive skill.

These findings hint at a general ability of the brain to acquire new skills via flexibly connecting regions into novel modules, thus adding layers of organization, without harming existing abilities. One could imagine other cognitive strategies that use neural circuits in novel ways to overcome the limitations of our default circuitry.

## Resilience of skill circuits

The Memory Champion’s functional regional specialization through training provides ideas for enhancing cognitive resilience in aging. Mnemonic techniques, particularly the Method of Loci, seem to leverage skills such as egocentric navigation and tracking event boundaries in stories, that are relatively more preserved in healthy aging^73,82^. The Method of Loci has shown promise in the aging population, with individuals able to achieve high memory performance using the technique and remaining proficient with it at six-year follow-up^83^. Major changes in routine and relocation can bring stress and anxiety to some older adults, linked to difficulty adapting and acquiring new landmarks^84,85^. Moreover, the context-association network has been shown to be a mediator between the default-mode network and the hippocampus, with amplitude of the context-association network to medial temporal lobe functional connectivity associated with better episodic memory performance in older adults^13^. This suggests the possibility that robust context-association network function might support stable memory function during aging. Thus, exploring the use of the Method of Loci to reinforce context-association network functioning is an exciting prospect for maintaining cognitive abilities and fostering a resilient brain.

## Skill it to make it stick

The wide gulf between our ability to remember random sequences of abstract symbols and our ability to navigate without getting lost, or to remember salient social interactions, hints at the skills that mattered most across our evolutionary history^86^. Archaeological evidence suggests that modern human showed heightened mobility and spatially extensive social networks compared to Neanderthals^86^. Humans can accomplish complex cognitive tasks that still elude artificial intelligence, with a minuscule general-purpose memory for abstract symbols. Our brain evolved for human-specific tasks, especially navigation, storytelling^87^ and skill learning. As it turns out, the best memory is goal oriented. In the age of supercomputers and AI, human rote memorization is already near-obsolete. Thus, it might be a great time to make teaching, learning and working more naturally human friendly, and memorable, by leaning into our strengths. While computer memory and AI are beating expert humans at most things, we still have much to discover about our own abilities as mobile, social, energy efficient agents^88,89^.

## Acknowledgement

We would like to extend our sincere gratitude to Nelson Dellis for his invaluable contribution and participation in this study. This work was supported by NIH grants NS140256 (E.M.G.), NS123345, NS098482 (B.P.K.),MH129616 (T.O.L.), T32DA007261 (S.R.K), MH096773 (N.U.F.D.), MH122066 (N.U.F.D., E.M.G.), MH121276 (N.U.F.D., E.M.G.), MH124567 (N.U.F.D., E.M.G.), NS129521 (N.U.F.D., E.M.G.), and NS088590 (N.U.F.D.); by the Taylor Family Foundation (T.O.L.); by the Intellectual and Developmental Disabilities Research Center (N.U.F.D.); by the Kiwanis Foundation (N.U.F.D.); by the Washington University Hope Center for Neurological Disorders (B.P.K. and N.U.F.D.); by Mallinckrodt Institute of Radiology pilot funding (N.U.F.D.); by the McDonnell Center for Systems Neuroscience and the Brain and Behavior Research Foundation (R.J.C.). Computations were performed using the facilities of the Washington University Research Computing and Informatics Facility, which were partially funded by NIH grants S10OD025200, 1S10RR022984-01A1 and 1S10OD018091-01.

## Method

### Participants

#### Memory Champion

The Memory Champion was 31 years of age at the first visit and 37 years of age at the second visit, and practiced memory sports starting in 2010, representing 5 years and 11 years of training for each visit. The Memory Champion is a six-time United States Memory Champion who holds several current and former U.S. memory records along with the Grand Master of Memory title.

#### MSC participants

The MSC is composed of 10 neurotypical, right-handed, adult participants (24-34 years of age at the time of testing; 5 females), who were recruited from the Washington University in St. Louis community. Participants completed 10 sessions of scanning that were 1.5 hour each, with all scan sessions completed within 7 weeks. The MSC dataset is publicly available (https://openneuro.org/datasets/ds000224). Details of the dataset and processing pipeline have been previously described ^15^.

### PFM MRI acquisition

#### 2015 Memory Champion and MSC protocol

Data collection and processing pipeline for the Memory Champion followed the MSC protocol acquired in 2013, for the 2015 visit. All participants were scanned on a Siemens TRIO 3T across 12 sessions (2 sessions of structural MRI scans + 10 sessions of functional MRI scans). Structural images included four T1-weighted scans (TE = 3.74ms, TR = 2400ms, TI = 1000ms, flip angle = 8°, 0.8mm isotropic voxels, 224 sagittal slices), four T2-weighted images (TE = 479ms, TR = 3200ms, 0.8mm isotropic voxels, 224 sagittal, 224), four MRA and eight MRV scans.

Functional images included 300 minutes of eyes-open resting-state fMRI BOLD data (10 sessions x 30min/session) using a gradient-echo EPI BOLD sequence (TE = 27ms, TR = 2.2s, flip angle = 90°, 4mm isotropic voxels, 36 axial slices). Gradient echo field map images (one per session) were acquired with the same parameters. See (Gordon et al., 2017) for more details.

#### 2021 Memory Champion, MSC02 and MSC06 protocol

Each participant performed multiple MRI acquisition sessions for functional imaging. All participants were scanned on a Siemens TRIO 3T MRI scanner. For each session, the protocol included a series of task fMRI and resting-state fMRI BOLD data using the CMRR multiband (Mb)-ME sequence^90,91^ with five echoes (14.2 ms, 38.93 ms, 63.66 ms, 88.39 ms, 113.12 ms; TR = 1.761 s, 2 mm resolution, MB factor = 6, IPAT = 2, flip angle = 68°). Data was acquired in the PA phase encoding direction. Additionally, spin echo field maps were acquired in both AP and PA directions (SE, single-band, 3 frames per run, TR = 8.0 s, TE = 66 ms, flip angle = 90°).

This resulted in approximately 8 hours of fMRI data: 105 minutes of eyes-open resting-state fMRI BOLD data and 377 minutes of task fMRI for the Memory Champion. Table 1 provides details on the number of available minutes per task per participant. Two tasks were tailored to the Memory Champion’s performance of the Method of Loci and were not performed by the controls. Additionally, the two localizer tasks were acquired to better understand the neural response of the Memory Champion’s Method of Loci performance.

The order of task fMRI acquisition was randomized in each session. The resting-state scan followed by the deck of cards memory task were performed last. The memorization of the card order was scored immediately after the task, outside of the scanner.

### fMRI Task

#### Stanford VPN Localizer

The Stanford VPN localizer task is a functional localizer experiment used to define category-selective cortical regions^92^. The selected categories were adult face, body, instrument, corridor (labeled place), and digit. The task was a block design presentation per category, with a 1-back task instruction, implemented using Psychopy (version available at https://github.com/VPNL/fLoc).

All subsequent tasks were built using JSpsych. The reading span task was adapted from the versions available in the Experiment Factory^93^. Each task and each task repetition within a subject had an independent event or block presentation order, with an optimized jittering for design efficiency specific to our 1.761 seconds TR fMRI sequence, defined using NeuroDesign^94^. All tasks were optimized for hemodynamic response function detection power. See Supplementary Figs. 7, 8, 11 and 12 for task schema.

#### Action localizer

The action localizer task used stimuli from the BML-MoVi dataset (Biomotion Lab MoVi: A large multi-purpose human motion and video dataset^95^). The task comprises two different types of blocks: action and non-action blocks, performed as a 1-back task on short video stimuli. In the action blocks, the stimuli were reconstructed 3D body shapes of 13 different subjects performing one of 20 actions, such as crawling or throwing a ball. In the non-action blocks, the same videos were transformed into a vine (back-and-forth repetition) of the first second of the action video, reproducing all identical visual characteristics of the original video with body motion perception but without the perception of complete action movement.

The task was composed of three blocks of 10 videos for each category. Videos were followed by a time-jittered fixation cross. 1-back accuracy feedback and repetition of the instructions were provided for 9 seconds between blocks. See Supplementary Fig. 12 for task schema

#### Reading span task

The reading span task is a working memory task consisting of a memorization phase, a proactive recall phase, and a testing phase, repeated eight times. Each repetition involves a set of words to remember, ranging from four to seven words. Words are presented one at a time for 1.25 seconds, followed by a time-jittered fixation cross and a sentence. Participants are asked to remember the word and then judge whether the subsequent sentence is sensical or not, repeating this process four to seven times. This is followed by a screen marked ‘???’ for 7 seconds during which participants have to proactively recall the list of words they were asked to remember.

Finally, four to seven words are presented one by one on the screen with a question mark, and participants have to judge whether each word was part of the list they had to remember. This is followed by a time-jittered fixation cross. A short screen reading ‘new set’ indicates the start of a new set of word to remember, beginning again with the word memorization-sentence judgment alternation for the new set. See Supplementary Fig. 8 for task schema.

#### Deck of cards memory task

The deck of cards memory task aimed to target the neural correlates of the Method of Loci strategy. It comprised multiple sections to contrast the different elements of the strategy. The task begins with the viewing of cards. Cards appeared at a random location on the screen and the participant judges whether the card appeared on the left or right side of the screen. Both real and fake cards (with made-up symbols) were used in this section and the following one. Seventy cards were presented for 1 second each, followed by a time-jittered fixation cross. This was followed by a paired-association mentalization section, in which cards appeared at three positions on the screen, representing person, action and object association. 72 cards (real or fake) were presented one by one and the Memory Champion was instructed to mentally visualize the associated content. Each card appeared for 2 seconds, spaced 0.5 seconds apart. Then an 8-second instruction screen informed the Memory Champion of the next section: deck of cards memorization. A deck of cards was presented 3 cards at a time, as the Memory Champion chunks the deck of cards order 3-by-3 to construct scenes with 3 elements (action, person, and object). Each triplet of cards was shown for 6 seconds, spaced 0.3 seconds apart. This section was followed by a 1-minute fixation cross, during which the Memory Champion proactively recalled the order of the deck of cards. The proactive recall section was followed by 9 cued recall and testing. A triplet of cards from the deck just memorized was shown and the Memory Champion had 8 seconds to recall the three cards following it in the deck order. Then, a triplet of cards was proposed as a solution, and the Memory Champion had 3 seconds to judge whether it was the correct triplet. This deck of cards memorization, proactive recall, cued recall, and testing sequence was done twice, with a new deck of cards order to memorize. The Memory Champion then came out of the scanner and sorted two physical decks of cards in the order of the two decks of cards just memorized. Once both decks were sorted, accuracy was evaluated by the experimental team. See Supplementary Fig. 7 for task schema

#### Pi retrieval task

As a simulation of a pi challenge, in which the Memory Champion would be asked to find the 5 digits following a given 5 digits sequence within the first 10,000 digits of pi, a task was built to capture the Method of Loci recall strategy. The encoding strategy is similar to the deck of cards method, where an action-person-object association is assigned to each digit, then represented in a scene. The set of 5 digits is separated into the first three digits and the last three digits (with the middle digit represented twice). This results in a scene with two people performing two actions with two objects stored in a memory loci. The pi retrieval task aims to leverage this already encoded large item to capture the recall strategy in more detail. The task is composed of two sections: a cued recall phase (5 minutes) and a proactive recall phase (10 minutes).

During the first 5 minutes of the task, 5-digit sequences within the first 10,000 digits of the number pi are presented. The presentation of each digit sequence has three parts: (1) showing the first three digits for 4 seconds, (2) showing the last three digits for 4 seconds, during which the Memory Champion is instructed to compose the scene associated with the digit triplet, and (3) showing the full 5-digit sequence for 6 seconds, during which the Memory Champion is instructed to localize the scene in his memory palace. This sequence is repeated 15 times for different sets of 5 digits. Success and failure of location recall was accessed outside the MRI. As the Memory Champion recently retrained the first 5,000 digits, successes and failures perfectly align with location before and after the half mark. All digit series were part of the 10,000 digits, no catch trial was involved, to avoid deception and doubt. The Memory Champion could succeed (search + found) or fail (search only) to retrieve the series of digits within the 6 seconds allocated. We specifically contrasted the neural activation during search and successful retrieval of the localization of a 5-digit sequence from the first 10,000 digits of pi to differentiate networks involved in the longer search process versus localization of the digits.

This is followed by the proactive recall phase: a 10-minute fixation cross period during which the Memory Champion is instructed to recall the 10-000 first digits of pi in order from the start. See Supplementary Fig. 11 for task schema

#### HCP datasets and processing

We used data from 887 participants aged 22-35 years (410 female) from the HCP 1200 participants release^47^, each with 60 minutes of resting-state fMRI. A custom Siemens SKYRA 3.0T MRI scanner and a custom 32-channel head matrix coil were used to obtain high-resolution T1-weighted (MP-RAGE, TR = 2.4 s, 0.7 mm³ voxels) and BOLD contrast-sensitive (gradient-echo EPI, multiband factor 8, TR = 0.72 s, 2 mm³ voxels) images from each participant. BOLD acquisition included a left-to-right or right-to-left phase encoding, performed for 15 minutes of resting state twice for each participant. Data was downloaded from https://db.humanconnectome.org and used the standard HCP preprocessing pipeline^96^, which includes similar steps to our PFM preprocessing pipeline. For more information on acquisition and processing, see ^47,72,96–98^.

### PFM MRI preprocessing

#### Structural

Cortical surfaces were generated according to the procedure described in ^25^. Each participant’s averaged T1-weighted image, after field inhomogeneity correction by FSL FAST ^99^, was processed through the FreeSurfer recon-all pipeline to anatomically segment and label the amygdala, accumbens, caudate, cerebellum, hippocampus, pallidum, putamen, and thalamus in both hemispheres. This segmentation was used to create subcortical masks and anatomical surfaces for transforming the functional data into grayordinates (CIFTI format). The surfaces were then registered into fs_LR_32k surface space using the Multi-modal Surface Matching algorithm ^72^. Volumes were also registered to MNI152 space. Percent regional volumes of the Memory Champion and MSC controls are available in Supplementary Fig. 13.

#### Caudate segmentation

The caudate was segmented into head and body segments using anterior commissure-posterior commissure (AC-Pc) alignment via ACPC Detect^100^. Coronal slices of the AC and PC were used to delimit the end of the head and the beginning of the body segments within the caudate structure, which was segmented by FreeSurfer. The tail segment, located posterior to the PC slices, was not represented for all participants, as the caudate tail is a thin and elongated structure that might require higher resolution for accurate capture in some individuals. Therefore, analyses were performed only for the head and body segments of the caudate.

#### Functional preprocessing

Functional preprocessing follows the previously described pipeline of the MSC dataset ^25^ and involved temporal interpolation to correct for differences in slice acquisition timing, rigid-body correction of head movements, correction for susceptibility inhomogeneity-related distortions using field maps, and alignment to MNI152 space. For multi-echo data, echoes were optimally combined using the T2*-based echo weighting combination method^101^. Processing was done using 4dfp tools (https://readthedocs.org/projects/4dfp/).

#### Signal to noise ratio mask

Signal-to-noise ratio (SNR) is calculated on the data before noise regression as the average signal overtime divided by variance of signal overtime for each run. We used a SNR mask to remove region of low signal for the analysis. In the 2015 dataset, we use the average mask defined with the MSC data to maintain comparability with previous analysis with the MSC dataset. For the 2021 dataset, we define a SNR mask for each individual, as region with SNR above 20.

#### Resting state fMRI preprocessing

Resting state fMRI data denoising involved the removal of high-motion frames (framewise displacement [FD] > 0.1 mm for the single echo data and FD > 0.16 mm after low-pass filtering at 0.1 Hz for the multi-echo data), band-pass filtering (0.005 to 0.1 Hz), and regression of nuisance time series, including head movement parameters, the global signal (averaged and derivative across all gray-matter voxels), and orthogonalized waveforms extracted from ventricles, white matter, and extracranial tissues. The resting state fMRI data was then projected to the cortical surface as the last step. The data were smoothed using a two-dimensional 6-mm full-width half-maximum (FWHM) smoothing kernel. Additionally, to mitigate the effects of signal contamination from nearby cortical areas in the subcortex, the average time series of all cortical vertices within 20 mm of a subcortical voxel was included as a regressor to clean the subcortical volume, similar to strategies used in previous work on subcortical functional connectivity^102,103^.

#### Task-fMRI preprocessing

Task fMRI data were low-pass filtered at 0.1 Hz. The task fMRI data was then projected to the cortical surface. Task analyses were performed in grayordinates, following the HCP pipeline. 6 Head-movement, ventricle and white matter time series were selected and included in the FSL-based GLM analysis as confounds. Frames with high head movement (same selection parameters as resting state) were identified and each was included as an individual column in the confound matrix of the GLM. Only the resulting cortical surfaces from the task analysis were smoothed using a two-dimensional 6-mm full-width half-maximum (FWHM) smoothing kernel.

### fMRI processing

#### FMRI Task analysis

Task block designs and event-related designs were modeled using a double gamma HRF in a GLM analysis with FSL FEAT ^32^. Analysis was conducted independently on surface and subcortical grayordinates, following the HCP pipeline steps (release v4.3). Second-level analysis across runs within participants was performed using FSL, and the resulting z-scores were used to compare effects. Only positive BOLD activity was considered and displayed.

#### Network individual boundaries (MSHBM)

To define individual-specific network boundaries, the multi-session hierarchical Bayesian model (MS-HBM) method^104^ was used for each participant and each dataset. This method differentiates intra-subject (within-subject) from inter-subject (between-subject) network variability, which helps with a robust estimation of individual network boundaries. An 18-functional network identity template (Default Mode, Context Association, Visual-Lateral, Visual-Dorso-ventral Stream, Visual Primary, Visual Middle Temporal, Frontoparietal, Dorsal Attention, Peri-motor, Language, Salience, Action-mode, Parietal Memory, Auditory, Motor-Hand, Motor-Face, Motor-Foot, Auditory and Somato-Cognitive-Action), derived from ^44^, was used to inform the final step of network labeling.

#### FC quantification

Grayordinates from regions of interest or networks were used to average BOLD fMRI activity into time series. These time series were then used to calculate either full brain correlation maps or to calculate correlations between networks and regions. For seed map functional connectivity, all resting-state fMRI runs were concatenated, while resting-state fMRI runs were kept separate for run-specific FC, which was then used for testing group differences. For all analyses comparing the PFM participants to the HCP population, the atlas template network borders were used to calculate the network FC maps. For all analyses comparing the Memory Champion to controls with Precision Functional Mapping data, individual-specific network borders were used for analysis.

#### Module analysis

Seed FC maps from the high centrality FC region of the Memory Champion were calculated from the concatenated resting-state. Spatial correlations between seed maps were calculated and used to perform hierarchical clustering using the Ward algorithm to order the regions. Tests of the significance of the similarity between the specific regions’ seed maps were performed against the correlation similarity of 1,000 random seed maps with matching sizes to each of the high centrality seed regions. Random seeds were generated using Moran randomization to account for the spatial constraints of the individual brain on the surface projection and the volume mask. A z-score and percent correlation beyond the random distribution is presented in Extended data Fig. 4 and shows larger similarities than random forming clusters. A principal component analysis was also performed across the high centrality regions seed maps to validate the signal separation across different components, replicating the three clusters observed with the principal component analysis.

#### FC testing in PFM dataset

Resting-state run-specific FC values were tested for differences between the Memory Champion and controls and corrected within datasets (2015, 2021). For each region (anatomically or functionally defined) of interest, networks with positive FC to that region in the Memory Champion were considered for testing using independent t-tests and included in the False Discovery Rate (FDR) correction. For full network-to-network testing, all positive network-to-network connections were tested and corrected for FDR. Similarly, for full subcortical-to-network testing, all positive subcortical-to-network connections were included in the FDR correction. Significance was determined at p=0.05 after correction.

### Specificity and centrality analysis with HCP data

#### Gaussian Noise FC normalization

To compare FC between our datasets, in which the distribution of values is skewed differently due to differences in fMRI sequence, the number of volumes available, and preprocessing choices, we performed a mixture modeling normalization of the FC. This method is an adaptation of a technique used in previous functional connectivity studies to compare FC^105–108^. The method aims to model the Gaussian noise independently of the FC signal from the tails of the distribution, in order to normalize the FC distribution by the Gaussian noise distribution. To achieve this, we aggregate all FC network maps for a subject to obtain a brain-wide fully defined FC distribution and perform a Gaussian-Gamma mixture model. The two-tailed gamma distributions capture the FC signal, while the main Gaussian captures the noise. The mean and standard deviation of the Gaussian distribution are then used to scale the FC distribution. This individual-specific rescaled FC allows a more accurate comparison of network FC maps across individuals. This scaling is done at the structure level (cortex, and each FreeSurfer-segmented subcortical structure from the subcortical mask) to better address signal differences between sequence and scanner, especially different signal quality in the subcortex. This rescaling is performed for each individual in the HCP dataset and the PFM datasets and results in Z score FC maps.

#### High centrality detection

Once the rescaled FC network maps are obtained for each individual, the rescaled network FC maps of the Memory Champion are compared to each map of the HCP participants. Quantification of the percentage of HCP FC maps below the FC values of the Memory Champion is performed at the grayordinate level. This results in a ratio describing how specifically strong the Memory Champion’s FC is locally for each network in our template. This quantification is performed for both visits of the Memory Champion. Regions with FC beyond 95% of controls are selected for further analysis. Regions selected for more than one network and larger than 20 grayordinates are defined as high centrality regions.

#### Neurosynth meta-analytic term map analysis

To characterize the functional role of specific regions, we explore the Neurosynth ^52^ terms maps, which are statistical maps grouping references of neural activation through automated literature review. We used NiMare^109^ to download the terms maps and removed all terms that were related to anatomy, functional network, disorders, and conjunction words. Each map was max scaled and the mean value of voxels from each of the high centrality Memory Champion regions and from the combined voxels of regions within a cortical module was calculated. The 15 terms with the highest values were selected and displayed in a wordcloud map with font size weighted by the term average value using the wordcloud python package.

#### Winner-take-all network representation in subcortex

To determine where networks are represented in the subcortex and to create a consensus map, a winner-take-all analysis is performed across average network maps for the HCP population and MSC participants. Average network maps are calculated from the rescaled FC maps to better adjust for variation in noise between subjects in each dataset. The network with the highest FC value is identified as the winner for a given grayordinate, resulting in maps with network labels for both the HCP population and MSC participants.

#### Network territory in subcortex

While the winner-take-all approach provides a condensed visual summary, it masks the richness of network representation under the winning network. To quantify the overlap of the specificity pattern with the network representation in the subcortex, we opted to quantify the overlap with network territories. To account for uncertainty in the selection of the winner, we used a Dempster-Shafer theoretic model, as presented in^110^, to weigh the representation of each network at each voxel and evaluate which networks are competitors to the winning network. This results in a map for each network, delineating regions where network connectivity is strongly represented and competing for the winner labeling, which we refer to as network territory.

#### Spatial Significant testing

To evaluate whether a result is significantly spatially represented in a cortical network or subcortical structure, results are tested against null distribution maps built using the Moran randomization strategy ^111^. This randomization is constrained by spatial distance in the data, enabling us to test both cortical and subcortical spatial results with the same algorithm. Spatial randomization can be built relative to each individual brain’s spatial constraints, which is highly appropriate for precision functional mapping analysis, as previously demonstrated in ^43^.

After building 1000 randomized maps that respect the distribution value of the result under evaluation, average values are calculated for non-thresholded maps per individual-specific network borders and subcortical structures. For thresholded maps, the percentage overlap of the result with network and subcortical structures is calculated. A p-value is then calculated by comparing the true value to the random maps distribution values. These p-values are corrected for multiple comparisons across networks and subcortical structures using FDR.

Testing is performed for network and high centrality significant representation in task contrast, FC maps and the Memory Champion memory networks specificity overlap with controls network territories in the subcortex. For the specificity to network territory testing, the overlap is calculated over the entire subcortical representation of the network territory, not by individual subcortical structure, to simplify the interpretation.

## Reference

1. Forsberg, A., Adams, E. J. & Cowan, N. Chapter One - The role of working memory in long-term learning: Implications for childhood development. in Psychology of Learning and Motivation (ed. Federmeier, K. D.) vol. 74 1–45 (Academic Press, 2021).

2. Miller, G. A. The magical number seven, plus or minus two: Some limits on our capacity for processing information. Psychological Review 63, 81–97 (1956).

3. Cowan, N. The Magical Mystery Four: How is Working Memory Capacity Limited, and Why? Curr Dir Psychol Sci 19, 51–57 (2010).

4. Dellis, N. Memory Superpowers!: An Adventurous Guide to Remembering What You Don’t Want To Forget. (Abrams Books for Young Readers, 2020).

5. Squire, L. R. The Legacy of Patient H.M. for Neuroscience. Neuron 61, 6–9 (2009).

6. Rolls, E. T. & Treves, A. A theory of hippocampal function: New developments. Progress in Neurobiology 238, 102636 (2024).

7. Henke, K. A model for memory systems based on processing modes rather than consciousness. Nat Rev Neurosci 11, 523–532 (2010).

8. Raichle, M. E. The brain’s default mode network. Annu Rev Neurosci 38, 433–447 (2015).

9. Gilmore, A. W., Kalinowski, S. E., Milleville, S. C., Gotts, S. J. & Martin, A. Identifying task-general effects of stimulus familiarity in the parietal memory network. Neuropsychologia 124, 31–43 (2019).

10. Wei, X. & Buckner, R. L. Old-New Recognition Memory Revisited. 2025.11.11.687844 Preprint at 10.1101/2025.11.11.687844 (2025).

11. Buzsáki, G. & Moser, E. I. Memory, navigation and theta rhythm in the hippocampal-entorhinal system. Nat Neurosci 16, 130–138 (2013).

12. Czajkowski, R. et al. Encoding and storage of spatial information in the retrosplenial cortex. Proceedings of the National Academy of Sciences 111, 8661–8666 (2014).

13. Kaboodvand, N., Bäckman, L., Nyberg, L. & Salami, A. The retrosplenial cortex: A memory gateway between the cortical default mode network and the medial temporal lobe. Hum Brain Mapp 39, 2020–2034 (2018).

14. Aminoff, E. M., Kveraga, K. & Bar, M. The role of the parahippocampal cortex in cognition. Trends Cogn Sci 17, 379–390 (2013).

15. Gordon, E. M. et al. Precision Functional Mapping of Individual Human Brains. Neuron 95, 791–807.e7 (2017).

16. Egner, T. The Caudate Nucleus Mediates Learning of Stimulus–Control State Associations. J Neurosci 37, 1028–1038 (2017).

17. Kwon, Y. H. et al. Situating the salience and parietal memory networks in the context of multiple parallel distributed networks using precision functional mapping. Cell Rep 44, 115207 (2025).

18. Wei, X. & Buckner, R. L. Old-New Recognition Memory Revisited. BioRxiv 2025.11.11.687844 (2025) doi:10.1101/2025.11.11.687844.

19. Seger, C. A. & Cincotta, C. M. The Roles of the Caudate Nucleus in Human Classification Learning. J Neurosci 25, 2941–2951 (2005).

20. Poldrack, R. A. & Packard, M. G. Competition among multiple memory systems: converging evidence from animal and human brain studies. Neuropsychologia 41, 245–251 (2003).

21. Hughes, J. E. A. et al. Savant syndrome has a distinct psychological profile in autism. Molecular Autism 9, 53 (2018).

22. Dresler, M. et al. Mnemonic Training Reshapes Brain Networks to Support Superior Memory. Neuron 93, 1227–1235.e6 (2017).

23. Roediger, H. L. The effectiveness of four mnemonics in ordering recall. Journal of Experimental Psychology: Human Learning and Memory 6, 558–567 (1980).

24. Dellis, N. Improve Your Memory: With the Astonishing Memory Palace Technique: Includes 52 Cards, 64-Page Book, and a Fold-Out Memory Map Poster. (Arcturus Publishing, 2024).

25. Laumann, T. O. et al. Functional System and Areal Organization of a Highly Sampled Individual Human Brain. Neuron 87, 657–670 (2015).

26. Gratton, C. et al. Functional Brain Networks Are Dominated by Stable Group and Individual Factors, Not Cognitive or Daily Variation. Neuron 98, 439–452.e5 (2018).

27. Wagner, I. C. et al. Durable memories and efficient neural coding through mnemonic training using the method of loci. Sci Adv 7, eabc7606 (2021).

28. Maguire, E. A., Valentine, E. R., Wilding, J. M. & Kapur, N. Routes to remembering: the brains behind superior memory. Nat Neurosci 6, 90–95 (2003).

29. Ren, J. et al. Method of loci training yields unique neural representations that support effective memory encoding. 2025.02.24.639840 Preprint at 10.1101/2025.02.24.639840 (2025).

30. D’Esposito, M. & Postle, B. R. The Cognitive Neuroscience of Working Memory. Annual Review of Psychology 66, 115–142 (2015).

31. Müller, N. C. J. et al. Hippocampal–caudate nucleus interactions support exceptional memory performance. Brain Struct Funct 223, 1379–1389 (2018).

32. Fox, K. & Stryker, M. Integrating Hebbian and homeostatic plasticity: introduction. Philosophical Transactions of the Royal Society B: Biological Sciences 372, 20160413 (2017).

33. Bassett, D. S. et al. Dynamic reconfiguration of human brain networks during learning. Proceedings of the National Academy of Sciences 108, 7641–7646 (2011).

34. Rubinov, M. & Sporns, O. Complex network measures of brain connectivity: Uses and interpretations. NeuroImage 52, 1059–1069 (2010).

35. Ooi, L. Q. R. et al. Longer scans boost prediction and cut costs in brain-wide association studies. Nature 1–10 (2025) doi:10.1038/s41586-025-09250-1.

36. Newbold, D. J. et al. Plasticity and Spontaneous Activity Pulses in Disused Human Brain Circuits. Neuron 107, 580–589.e6 (2020).

37. Pritschet, L. et al. Neuroanatomical changes observed over the course of a human pregnancy. Nat Neurosci 27, 2253–2260 (2024).

38. Krimmel, S. R. et al. The human brainstem’s red nucleus was upgraded to support goal-directed action. Nat Commun 16, 3398 (2025).

39. Marek, S. et al. Spatial and Temporal Organization of the Individual Human Cerebellum. Neuron 100, 977–993.e7 (2018).

40. Badke D’Andrea, C., et al. Action-mode subnetworks for decision-making, action control, and feedback. Proceedings of the National Academy of Sciences 122, e2502021122 (2025).

41. Gordon, E. M. et al. A somato-cognitive action network alternates with effector regions in motor cortex. Nature 617, 351–359 (2023).

42. Braga, R. M. & Buckner, R. L. Parallel Interdigitated Distributed Networks within the Individual Estimated by Intrinsic Functional Connectivity. Neuron 95, 457–471.e5 (2017).

43. Chauvin, R. J. et al. Disuse-driven plasticity in the human thalamus and putamen. Cell Reports 44, 115570 (2025).

44. Lynch, C. J. et al. Expansion of a frontostriatal salience network in individuals with depression. 2023.08.09.551651 Preprint at 10.1101/2023.08.09.551651 (2023).

45. Laumann, T. O. et al. Brain network reorganisation in an adolescent after bilateral perinatal strokes. The Lancet Neurology 20, 255–256 (2021).

46. Nahas, Z., et al. Personalized Adaptive Cortical Electro-stimulation (PACE) in Treatment-Resistant Depression. Preprint at 10.31234/osf.io/5c3ba_v1 (2025).

47. Van Essen, D. C. et al. The Human Connectome Project: A data acquisition perspective. NeuroImage 62, 2222–2231 (2012).

48. Hodgson, V. J., Lambon Ralph, M. A. & Jackson, R. L. Multiple dimensions underlying the functional organization of the language network. NeuroImage 241, 118444 (2021).

49. Vann, S. D., Aggleton, J. P. & Maguire, E. A. What does the retrosplenial cortex do? Nat Rev Neurosci 10, 792–802 (2009).

50. Delikishkina, E., Lingnau, A. & Miceli, G. Neural correlates of object and action naming practice. Cortex 131, 87–102 (2020).

51. Dosenbach, N. U. F., Raichle, M. & Gordon, E. M. The brain’s cingulo-opercular action-mode network. https://doi.org/10.31234/osf.io/2vt79 (2024) doi:10.31234/osf.io/2vt79.

52. Yarkoni, T., Poldrack, R. A., Nichols, T. E., Van Essen, D. C. & Wager, T. D. Large-scale automated synthesis of human functional neuroimaging data. Nat Methods 8, 665–670 (2011).

53. Thommesen, K. K. et al. Neural underpinnings of memory encoding and retrieval: Validation of a novel ecologically valid fMRI paradigm. Neuroscience Applied 3, 104084 (2024).

54. Kim, H. Neural activity that predicts subsequent memory and forgetting: a meta-analysis of 74 fMRI studies. Neuroimage 54, 2446–2461 (2011).

55. Grön, G. et al. Hippocampal Activations during Repetitive Learning and Recall of Geometric Patterns. Learn Mem 8, 336–345 (2001).

56. Delazer, M. et al. Learning by strategies and learning by drill—evidence from an fMRI study. NeuroImage 25, 838–849 (2005).

57. Borders, A. A., Aly, M., Parks, C. M. & Yonelinas, A. P. The hippocampus is particularly important for building associations across stimulus domains. Neuropsychologia 99, 335–342 (2017).

58. Meltzer, J. A. & Constable, R. T. Activation of human hippocampal formation reflects success in both encoding and cued recall of paired associates. Neuroimage 24, 384–397 (2005).

59. Clark, I. A., Kim, M. & Maguire, E. A. Verbal Paired Associates and the Hippocampus: The Role of Scenes. J Cogn Neurosci 30, 1821–1845 (2018).

60. Seger, C. A. & Miller, E. K. Category Learning in the Brain. Annual review of neuroscience 33, 203 (2010).

61. Williams, Z. M. & Eskandar, E. N. Selective enhancement of associative learning by microstimulation of the anterior caudate. Nat Neurosci 9, 562–568 (2006).

62. Scimeca, J. M. & Badre, D. Striatal contributions to declarative memory retrieval. Neuron 75, 380–392 (2012).

63. Seger, C. A. & Cincotta, C. M. Dynamics of Frontal, Striatal, and Hippocampal Systems during Rule Learning. Cereb Cortex 16, 1546–1555 (2006).

64. Colombo, D. et al. Egocentric and allocentric spatial reference frames in aging: A systematic review. Neuroscience & Biobehavioral Reviews 80, 605–621 (2017).

65. Roediger, H. L. & Karpicke, J. D. Test-Enhanced Learning: Taking Memory Tests Improves Long-Term Retention. Psychol Sci 17, 249–255 (2006).

66. Gupta, M. W. & Rickard, T. C. The effects of declarative learning on early and late motor skill learning. npj Sci. Learn. 11, 5 (2025).

67. Doyon, J. et al. Contributions of the basal ganglia and functionally related brain structures to motor learning. Behavioural Brain Research 199, 61–75 (2009).

68. Haith, A. M. & Krakauer, J. W. The multiple effects of practice: skill, habit and reduced cognitive load. Current Opinion in Behavioral Sciences 20, 196–201 (2018).

69. Farrokhi, A., Habibi, M. & Daliri, M. R. Exploring the Impact of Declarative Learning on the Consolidation of Acquired Motor Skills Under Valence Feedback. Hum Brain Mapp 46, e70105 (2025).

70. Smallwood, J. et al. The default mode network in cognition: a topographical perspective. Nat Rev Neurosci 22, 503–513 (2021).

71. Tu, J. C. et al. Commonality and Variability in Functional Networks In Young Children Under 5 Years Old. 2025.09.12.675913 Preprint at 10.1101/2025.09.12.675913 (2025).

72. Glasser, M. F. et al. A multi-modal parcellation of human cerebral cortex. Nature 536, 171–178 (2016).

73. Reagh, Z. M., Delarazan, A. I., Garber, A. & Ranganath, C. Aging alters neural activity at event boundaries in the hippocampus and Posterior Medial network. Nat Commun 11, 3980 (2020).

74. Roediger III, H. L. & Zaromb, F. M. Memory for actions: How different? in Memory, aging and the brain: A Festschrift in honour of Lars-Göran Nilsson 24–52 (Psychology Press, New York, NY, US, 2010).

75. Kullmann, W. Homer and Historical Memory. in Signs of Orality 95–113 (Brill, 1999). doi:10.1163/9789004351424_007.

76. Neale, M. Songlines: The Power and Promise. (Thames & Hudson Australia, Port Melbourne, Victoria, 2020).

77. Mar, R. A., Li, J., Nguyen, A. T. P. & Ta, C. P. Memory and comprehension of narrative versus expository texts: A meta-analysis. Psychon Bull Rev 28, 732–749 (2021).

78. Goldstein, A. G., Chance, J. E., Hoisington, M. & Buescher, K. Recognition memory for pictures: Dynamic vs. static stimuli. Bulletin of the Psychonomic Society 20, 37–40 (1982).

79. Antony, J., Lozano, A., Dhoat, P., Chen, J. & Bennion, K. Causal and Chronological Relationships Predict Memory Organization for Nonlinear Narratives. J Cogn Neurosci 36, 2368–2385 (2024).

80. Schlaggar, B. L. & McCandliss, B. D. Development of Neural Systems for Reading. Annual Review of Neuroscience 30, 475–503 (2007).

81. Marano, G. et al. The Neuroscience Behind Writing: Handwriting vs. Typing—Who Wins the Battle? Life (Basel) 15, 345 (2025).

82. Colombo, D. et al. Egocentric and allocentric spatial reference frames in aging: A systematic review. Neuroscience & Biobehavioral Reviews 80, 605–621 (2017).

83. Gross, A. L. et al. Do older adults use the Method of Loci? Results from the ACTIVE Study. Exp Aging Res 40, 140–163 (2014).

84. Fealy, S. et al. Psychological interventions designed to reduce relocation stress for older people transitioning into permanent residential aged care: a systematic scoping review. Aging & Mental Health 28, 1197–1208 (2024).

85. West, G. L. et al. Landmark-dependent Navigation Strategy Declines across the Human Life-Span: Evidence from Over 37,000 Participants. J Cogn Neurosci 35, 452–467 (2023).

86. Burke, A. Spatial abilities, cognition and the pattern of Neanderthal and modern human dispersals. Quaternary International 247, 230–235 (2012).

87. Smith, D. et al. Cooperation and the evolution of hunter-gatherer storytelling. Nat Commun 8, 1853 (2017).

88. Shevlin, H., Vold, K., Crosby, M. & Halina, M. The limits of machine intelligence. EMBO Rep 20, e49177 (2019).

89. Korteling, J. E. (Hans)., van de Boer-Visschedijk, G. C., Blankendaal, R. A. M., Boonekamp, R. C. & Eikelboom, A. R. Human-versus Artificial Intelligence. Front Artif Intell 4, 622364 (2021).

90. Feinberg, D. A. et al. Multiplexed Echo Planar Imaging for Sub-Second Whole Brain FMRI and Fast Diffusion Imaging. PLOS ONE 5, e15710 (2010).

91. Moeller, S. et al. Multiband multislice GE-EPI at 7 tesla, with 16-fold acceleration using partial parallel imaging with application to high spatial and temporal whole-brain fMRI. Magn Reson Med 63, 1144–1153 (2010).

92. Stigliani, A., Weiner, K. S. & Grill-Spector, K. Temporal Processing Capacity in High-Level Visual Cortex Is Domain Specific. J Neurosci 35, 12412–12424 (2015).

93. Sochat, V. The Experiment Factory: Reproducible Experiment Containers. Journal of Open Source Software 3, 521 (2018).

94. Durnez, J., Blair, R. & Poldrack, R. Neurodesign: Optimal experimental designs for task fMRI. bioRxiv 119594 (2018) doi:10.1101/119594.

95. Ghorbani, S. et al. MoVi: A large multi-purpose human motion and video dataset. PLOS ONE 16, e0253157 (2021).

96. Glasser, M. F. et al. The Minimal Preprocessing Pipelines for the Human Connectome Project. Neuroimage 80, 105–124 (2013).

97. Smith, S. M. et al. Resting-state fMRI in the Human Connectome Project. Neuroimage 80, 144–168 (2013).

98. Glasser, M. F. et al. The Human Connectome Project’s neuroimaging approach. Nat Neurosci 19, 1175–1187 (2016).

99. Jenkinson, M., Beckmann, C. F., Behrens, T. E. J., Woolrich, M. W. & Smith, S. M. FSL. Neuroimage 62, 782–790 (2012).

100. Ardekani, B. A. & Bachman, A. H. Model-based automatic detection of the anterior and posterior commissures on MRI scans. Neuroimage 46, 677–682 (2009).

101. Posse, S. et al. Enhancement of BOLD-contrast sensitivity by single-shot multi-echo functional MR imaging. Magn Reson Med 42, 87–97 (1999).

102. Greene, D. J. et al. Developmental Changes in the Organization of Functional Connections between the Basal Ganglia and Cerebral Cortex. J. Neurosci. 34, 5842–5854 (2014).

103. Choi, E. Y., Yeo, B. T. T. & Buckner, R. L. The organization of the human striatum estimated by intrinsic functional connectivity. J Neurophysiol 108, 2242–2263 (2012).

104. Kong, R. et al. Spatial Topography of Individual-Specific Cortical Networks Predicts Human Cognition, Personality, and Emotion. Cereb Cortex 29, 2533–2551 (2019).

105. Bielczyk, N. Z. et al. Thresholding functional connectomes by means of mixture modeling. NeuroImage https://doi.org/10.1016/j.neuroimage.2018.01.003 (2018) doi:10.1016/j.neuroimage.2018.01.003.

106. Chauvin, R. J., Mennes, M., Buitelaar, J. K. & Beckmann, C. F. Assessing age-dependent multi-task functional co-activation changes using measures of task-potency. Dev Cogn Neurosci https://doi.org/10.1016/j.dcn.2017.11.011 (2017) doi:10.1016/j.dcn.2017.11.011.

107. Chauvin, R. J., Mennes, M., Llera, A., Buitelaar, J. K. & Beckmann, C. F. Disentangling common from specific processing across tasks using task potency. NeuroImage 184, 632–645 (2019).

108. Chauvin, R. J. et al. Task-generic and task-specific connectivity modulations in the ADHD brain: an integrated analysis across multiple tasks. Transl Psychiatry 11, 1–10 (2021).

109. Salo, T. Developing and Validating Open Source Tools for Advanced Neuroimaging Research. FIU Electronic Theses and Dissertations https://digitalcommons.fiu.edu/etd/5010 (2022).

110. Napoli, N. J., Barnes, L. E., Premaratne, K. & IEEE. Correlation Coefficient Based Template Matching: Accounting for Uncertainty in Selecting the Winner. 2015 18TH INTERNATIONAL CONFERENCE ON INFORMATION FUSION (FUSION) 311–318 (2015).

111. Markello, R. D. et al. neuromaps: structural and functional interpretation of brain maps. Nat Methods 1–8 (2022) doi:10.1038/s41592-022-01625-w.

