## Extended Data Figures for "Brain organization of a memory champion"

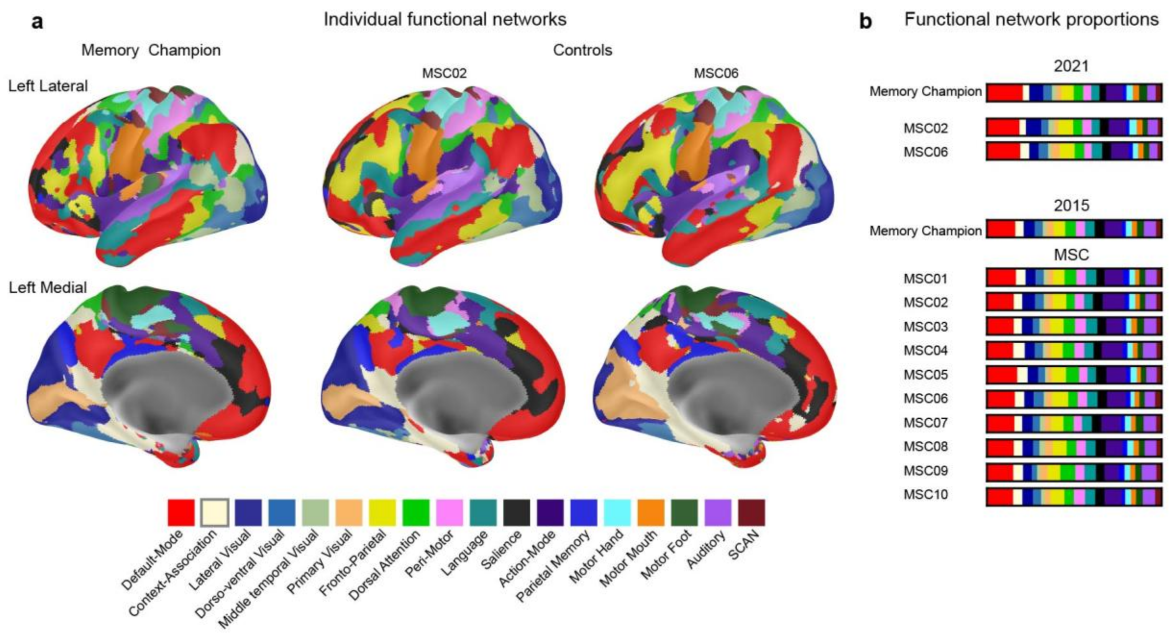


**Extended Data Figure 1: Individual-specific functional networks in PFM participants. a)** Individual-specific functional networks in the 2021 dataset. Left inflated lateral (top) and medial (bottom) views of the individual network labeling for the Memory Champion (left) and the two controls (MSC02 center, MSC06 right). **b)** Proportion of cortical representation of each network, comparing the Memory Champion to MSC02 and MSC06 in the 2021 data and the Memory Champion 2015 data to the MSC dataset. 2015 and MSC individual network labeling are available in Supplementary Fig. 1.

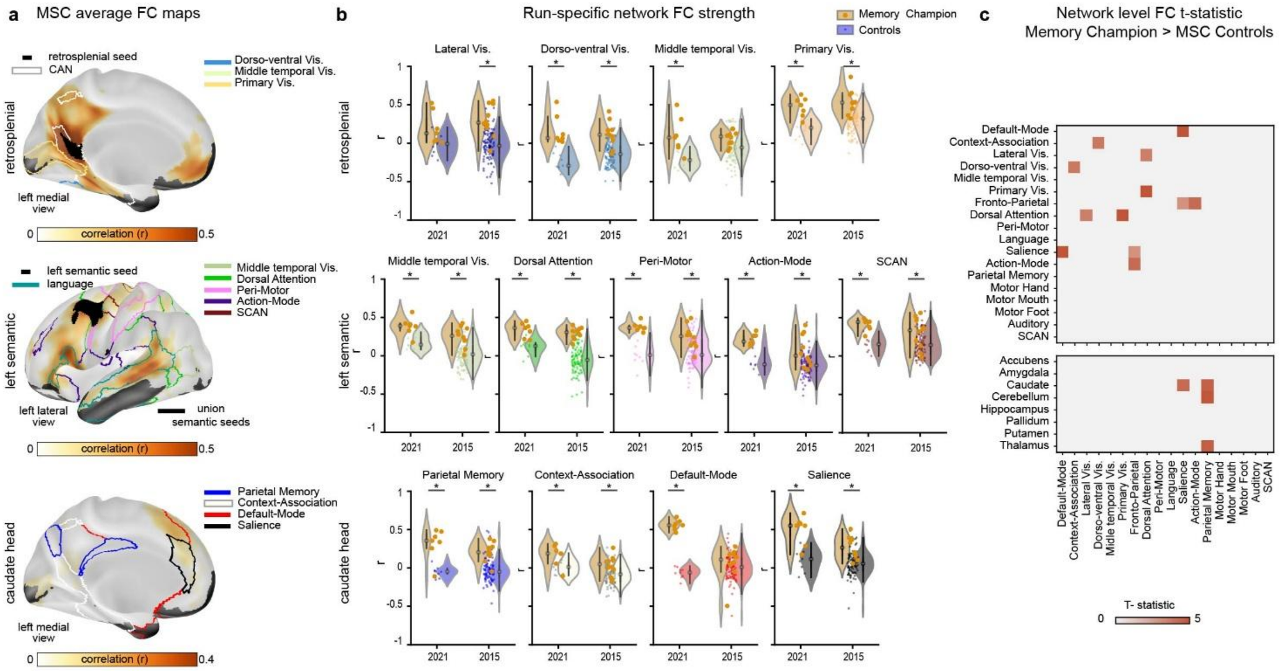


**Extended Data Figure 2: Comparison of functional connectivity in the Memory Champion to repeatedly sampled controls. a)** Average whole-brain FC maps for representative brain regions identified in Fig. 1c for the MSC controls. The top 20% of the strongest connections are shown. The top row shows correlation (r) for the retrosplenial cortex (Brodmann areas 26, 29, and 30, part of the contextual association network -white border), the middle row shows the left semantic region (area 55b; language network -teal border), and the bottom row shows the head of the left caudate. Full views are available in Supplementary Fig. 3. **b)** Distribution of FC values for each rest run in the Memory Champion and PFM controls. FC values between regions of panel A and individualized networks for each rest run were independently tested for significant differences with controls (controls are displayed in the color of the network, and the Memory Champion in orange). Only violin plots for networks with significant differences in both the 2015 and 2021 datasets are displayed, and additional networks that have comparative values : other visual networks for the retrosplenial and other memory networks for the caudate head. * indicates significance at $P < 0.05$ after FDR correction across connected networks. **c)** Average T-statistics for 2015 and 2021 differences in FC between the Memory Champion and controls for all significant increased FC, when testing all network-by-network and structure testing (one-tailed independent t-test, $P < 0.05$ FDR corrected). SCAN stands for somato-cognitive action network.

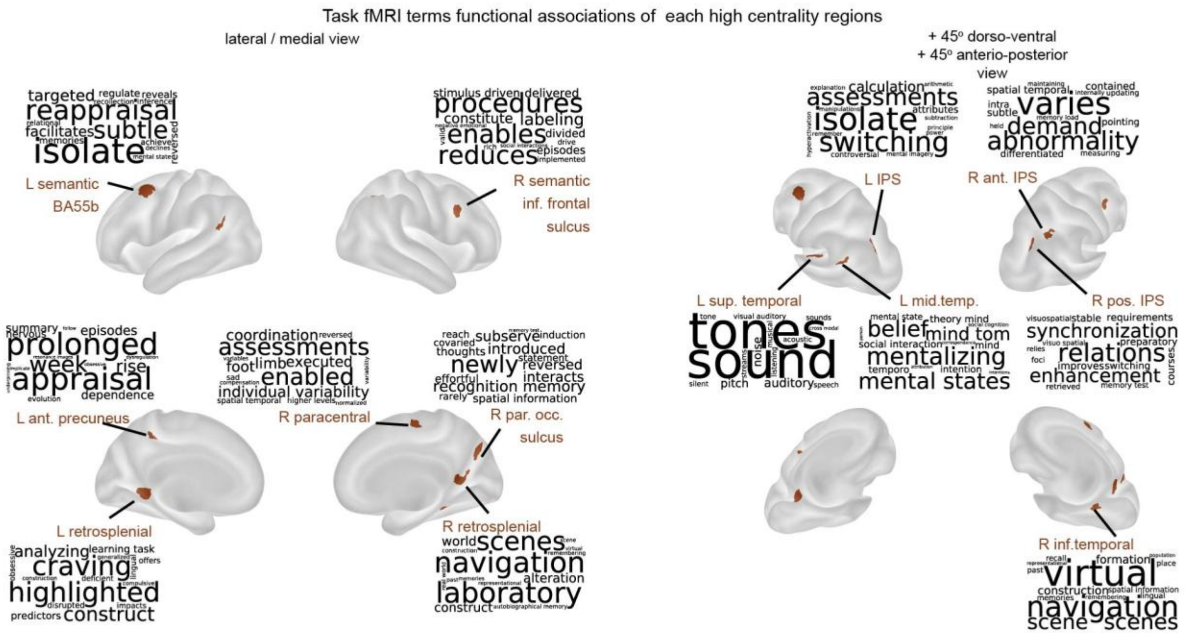


**Extended Data Figure 3: Top 15 meta-analytic data base terms (Neurosynth) associated with high centrality regions**. Inflated cortical views (lateral and medial on the left, rotated on the right) of the high centrality regions of the Memory Champion compared to controls from Fig. 1c. Abbreviations stand for Left (L), Right (R), anterior (ant), posterior (pos), parietal occipital sulcus (Par. Occ. S), intraparietal sulcus (IPS), inferior (inf), middle (mid), superior (sup), and Brodmann Area (BA). Next to each of the Memory Champion's high centrality regions is a word cloud of the 15 Neurosynth terms associated with each region. The font size of the word matches the relative weight of the term among the top 15.

| name | Centroid MNI coordinate (x, y, z) | Grayordinate count | Network FC > 95% of HCP population | Module assignment |
|---|---|---|---|---|
| Right caudate | 12, 14, 6 | 121 | Parietal Memory, Salience, DMN, CAN, FPN, AMN, Dorsal Attention | subcortical |
| Left caudate | -13, 10, 10 | 95 | Parietal Memory, Salience, CAN, DMN, FPN, Dorsal Attention, AMN | subcortical |
| Cerebellum | 14, -80, -44 | 51 | SCAN, Peri-motor, Auditory, Motor Mouth, Motor Hand, Temporal middle Visual, AMN | subcortical |
| Right anterior intra parietal sulcus | 32, -49, 39 | 103 | SCAN, Peri-motor, Motor Mouth, Motor Hand, Temporal middle Visual, Dorsal Attention | Semantic |
| Left intra parietal sulcus | -50, -26, 10 | 69 | Peri-motor, Language, Temporal middle Visual, Dorso-ventral Visual, Motor Hand, Visual Lateral | Semantic |
| Right parieto-occipital sulcus | 18, -70, 29 | 109 | Motor Mouth, Motor Hand, Peri-motor, Visual Lateral, Dorso-ventral Visual, Primary Visual, Motor Foot, Temporal middle Visual, Auditory, Language, SCAN | Scene |
| Left middle temporal | -49, -53, 21 | 69 | Visual Lateral, Dorso-ventral Visual, Auditory, Temporal middle Visual, Primary Visual, Dorsal Attention | Scene |
| Right inferior temporal | 23, -42, -12 | 73 | Primary Visual, Visual lateral, Dorso-ventral Visual, Temporal middle Visual, AMN | Scene |

| | | | | |
|---|---|---|---|---|
| Left retro splenial | -15, -48, -2 | 146 | Visual lateral, Auditory, Dorso-ventral Visual, Dorsal Attention, SCAN, Peri-motor, Motor Hand, Primary Visual, Motor Mouth, Temporal middle Visual, AMN | Scene |
| Right posterior intra parietal sulcus | 34, -75, 36 | 115 | Primary Visual, Language, SCAN, Temporal middle Visual, Visual lateral, Motor Mouth | Scene |
| Right retrosplenial | 12, -53, 10 | 183 | Visual Lateral, Primary Visual, Dorso-ventral Visual, Temporal middle Visual, Auditory, Language | Scene |
| Left superior temporal | -28, -73, 39 | 92 | Primary Visual, Visual Lateral, Temporal middle Visual, Dorso-ventral Visual, SCAN | Semantic |
| Left anterior precuneus | -6, -41, 51 | 73 | Motor Hand, SCAN, Motor Mouth, Language, Auditory, Motor Foot | Semantic |
| Left semantic BA 55b | -42, 10, 50 | 233 | SCAN, Dorsal Attention, Peri-motor, Auditory, AMN, Dorso-ventral Visual, Temporal middle Visual, Visual Lateral, Motor Hand, Primary Visual | Semantic |
| Right semantic Inferior frontal sulcus | 38, 9, 36 | 93 | SCAN, Motor Hand, Auditory, Peri-motor, Motor Mouth, Motor Foot, Language | Semantic |
| Right paracentral | 2, -9, 52 | 78 | Peri-motor, Motor Hand, Auditory, SCAN | Semantic |

**Table E1. Regions of high centrality in the Memory Champion.** Regions larger than 20 voxels with more than 2 networks with FC stronger than 95% of the HCP population in both the 2015 and 2021 datasets of the Memory Champion, along with their centroid coordinates in MNI, grayordinate count, network list showing >95% population strength in % beyond strength order, and module assignment.

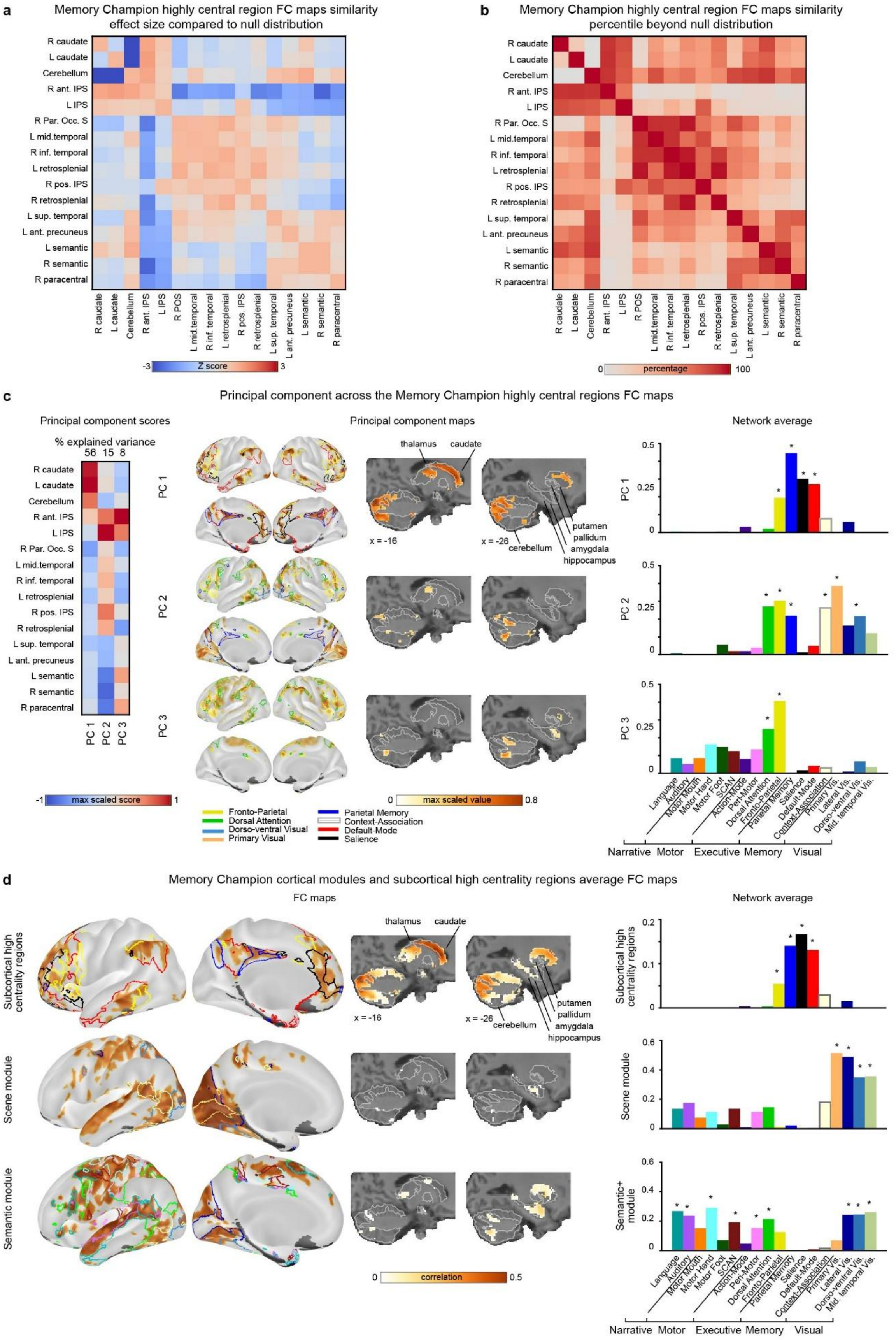
a
Memory Champion highly central region FC maps similarity
effect size compared to null distribution
R caudate
L caudate
Cerebellum
R ant. IPS
L IPS
R Par. Occ. S
L mid.temporal
R inf. temporal
L retrosplenial
R pos. IPS
R retrosplenial
L sup. temporal
L ant. precuneus
L semantic
R semantic
R paracentral
R POS
-3
Z score
3
b
Memory Champion highly central region FC maps similarity
percentile beyond null distribution
0
percentage
100
c
Principal component across the Memory Champion highly central regions FC maps
Principal component scores
% explained variance
56 15 8
PC 1
PC 2
PC 3
-1
max scaled score
1
Principal component maps
thalamus
caudate
putamen
pallidum
amygdala
hippocampus
cerebellum
x = -16
x = -26
0
max scaled value
0.8
Fronto-Parietal
Dorsal Attention
Dorso-ventral Visual
Primary Visual
Parietal Memory
Context-Association
Default-Mode
Salience
Network average
0.5
0.25
0
Language
Auditory
Motor Mouth
Motor Hand
Motor Foot
SCAN
Action-Mode
Peri-Motor
Dorsal Attention
Fronto-Parietal
Parietal Memory
Salience
Default-Mode
Context-Association
Primary Vis.
Lateral Vis.
Dorso-ventral Vis.
Mid. temporal Vis.
Narrative
Motor
Executive
Memory
Visual
d
Memory Champion cortical modules and subcortical high centrality regions average FC maps
FC maps
Subcortical high centrality regions
Scene module
Semantic module
0
correlation
0.5
Network average
0.2
0.1
0.6
0.4
Semantic+ module

**Extended Data Figure 4: Memory Champion high centrality regions module analyses. a)** Effect size against null of the resting-state Memory Champion high centrality regions FC maps similarly. Effect size is estimated as a z-score of the high centrality regions paired-seed map correlation compared to the null distribution of size-matching random paired-seed map correlation values. **b)** Percentile ranking of seed map similarity against the null distribution. An alternative metric to quantify the strength of functional similarity is to quantify the percentile ranking of the true value in the null distribution used in panel A. **c)** First three principal components across high centrality regions resting-state seed maps. Max-scaled principal component scores across high centrality regions are displayed in the left matrix. The top 20th percentile of each of the first three components (max-scaled) is displayed on the inflated brain and on x=-16 and x=-26 sagittal slices overlaid on the Memory Athlete's T1 (center). White borders outline the Freesurfer segmented anatomical structures. The average network values from the max-scaled positive cortical map of the first three components are displayed in the bar plot on the right, in the network color code. We note that the IPS regions show a particular behavior of sharing FC with all three first components (c), therefore their seed map similarity is ambiguous (a, b). We associate them with the semantic module as their strongest increase in centrality was with executive/motor networks rather than visual. **d)** Seed connectivity maps and network average for the Memory Champion's group of high centrality regions emerging from Fig. 3. The top 20th percentile of FC maps of the combined voxels across the cortical regions from the two modules as defined in Fig. 3, i.e., scene and semantic (top and middle lines, respectively) and three subcortical high centrality regions (left and right caudate and cerebellum) (bottom line) are displayed on left lateral and medial inflated cortical views (left) and in the subcortex with coronal slices at x = -16 and x = -26 in MNI coordinates. The bar plots (right) show the average positive connectivity of the seed FC map per functional network. * indicates significance tested against null map distribution and corrected for multiple comparisons (*P* < 0.05 after correction). Significant network borders are displayed on the inflated brains for each module.

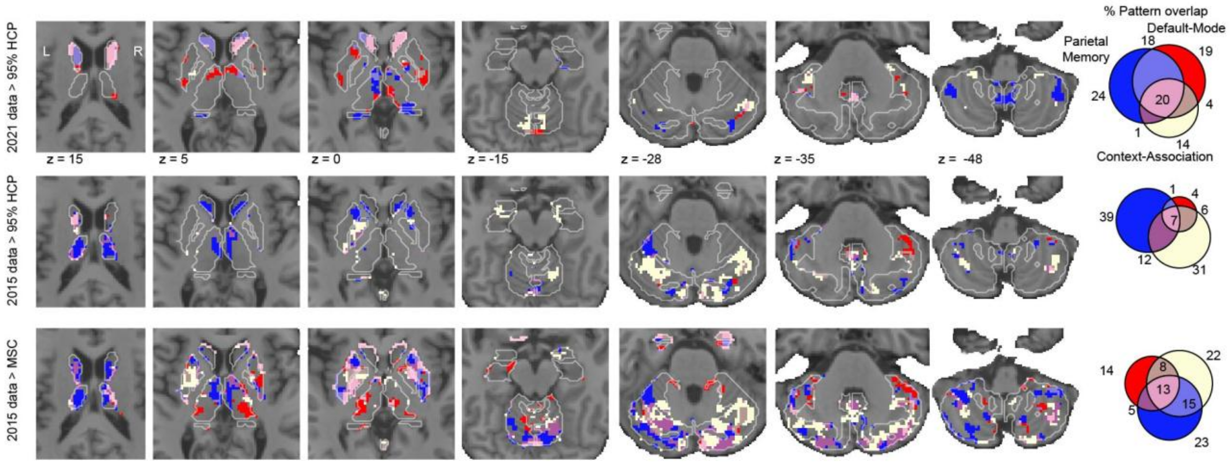


**Extended Data Figure 5: Functional connectivity between memory networks and subcortex in the Memory Champion.** Summary overlap of the three memory networks: parietal memory, default-mode, and context-association networks pattern of FC strength >95% of controls. The percentage of memory network overlapping and color code of the three memory networks are displayed on the Venn diagram (right). Results are displayed on the Memory

Champion T1 image. The first and second lines use the HCP population as controls for the 2021 and 2015 Memory Champion resting-state FC, respectively. The third line uses the MSC datasets as controls with the 2015 Memory Champion resting-state FC as replication as they share the same protocol.

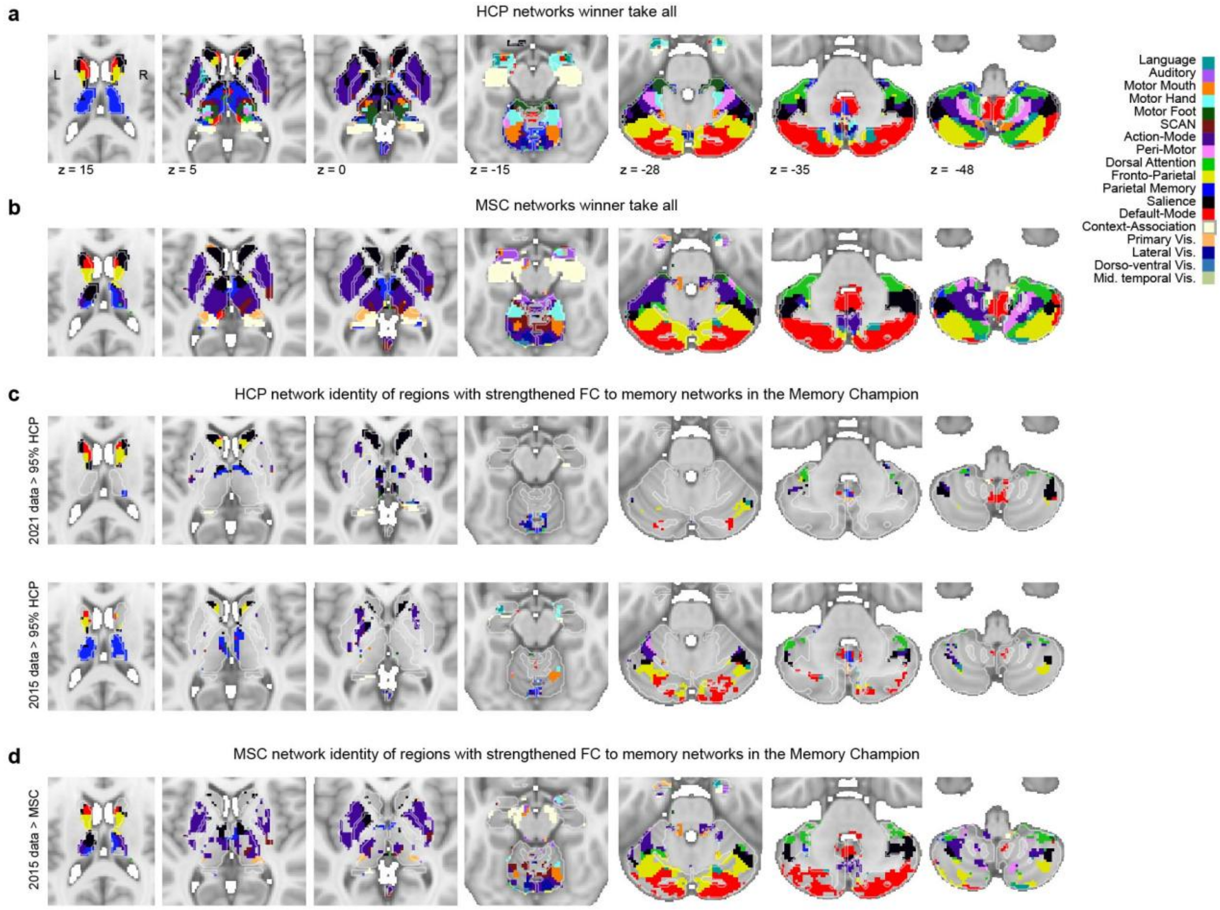


**Extended Data Figure 6: Parcellations of subcortex (winner-take-all) in controls and overlap with Memory Champion increased memory networks FC.** Average network winner-take-all (WTA) in subcortex for **a)** HCP and **b)** MSC data. Network identity, based on control data, of the regions with greater FC to memory networks in the Memory Champion for **c)** HCP and **d)** MSC data. The network winner-take-all (WTA) is shown from control data masked by greater memory network FC strength in the Memory Champion (>95% of HCP controls; > 100% MSC controls; see Extended Data Fig. 5). Results are displayed on the MNI average T1.

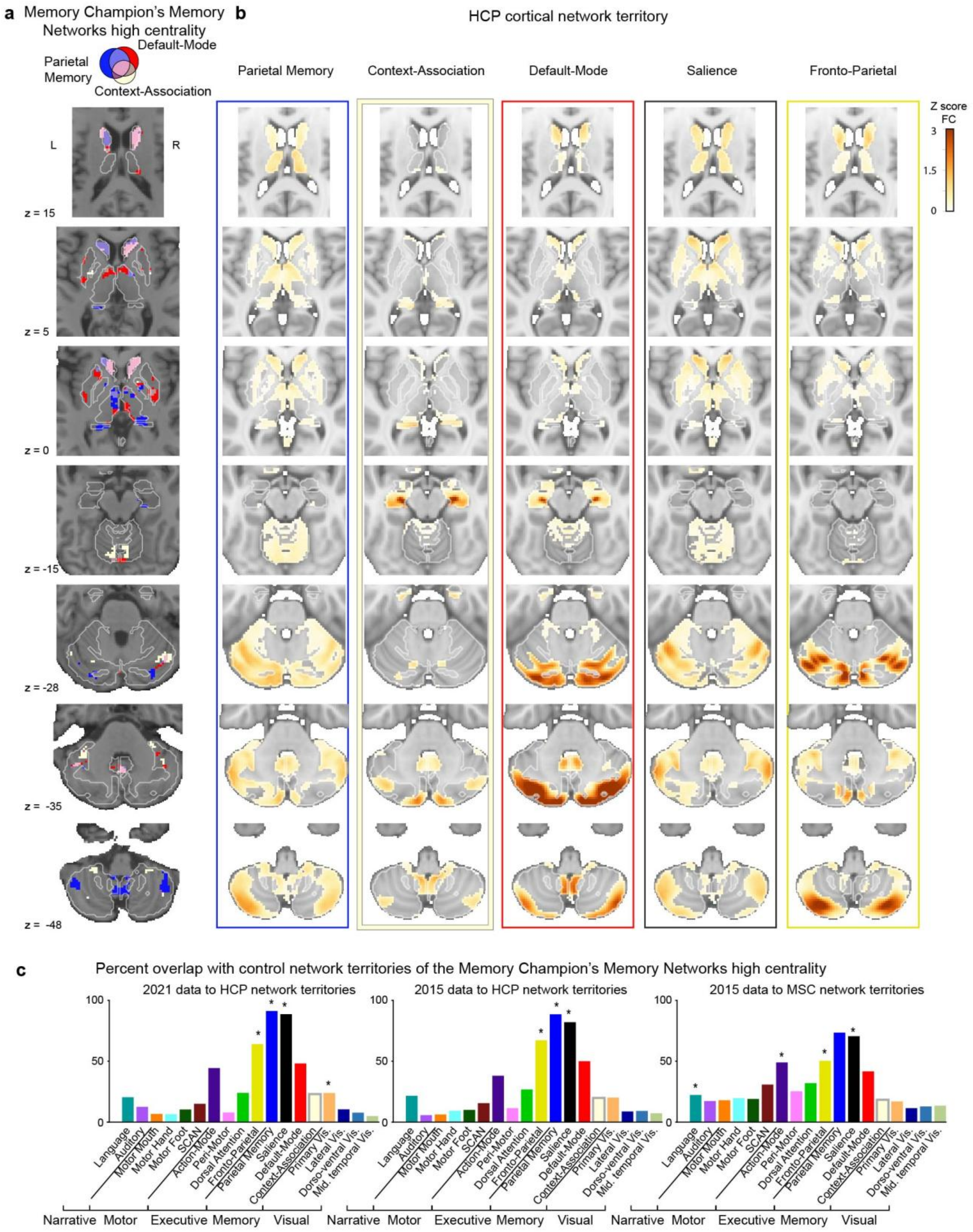


**Extended Data Figure 7: Quantification of Memory Champion's memory networks FC expansion in subcortex. a)** Pattern of increased FC between memory networks and subcortex. Combined memory networks with Memory champion 2021 data FC >95% of the HCP population. See the Venn diagram (top) for the network color code. Results are displayed on the Memory Champion T1 image. **b)** HCP average network territory in subcortex. Horizontal subcortical slices show (left to right column) Parietal Memory, Context-Association, Default-Mode, Salience, and Fronto-Parietal network FC strength z-score for subcortex. A network territory shows where a network is a competitor for winner-take-all labeling according to the Dempster-Shafer theoretical model (see Methods). Results are displayed on the MNI average T1. See Memory Champion network territories in Supplementary Fig. 6. **c)** Percentage of overlap with each controls network territories of the Memory Champion's memory network high centrality pattern. Comparison of the combined memory networks FC subcortical pattern >95% of controls and the controls network territories. Memory networks high centrality is defined in the Memory Champion 2021 resting state (left plot) and from 2015 resting state (middle and right plots) and compared to the HCP network territories (left and middle plots) and the 2015 MSC average network territories (right plot). Bar color code follows the network color code. * indicates significant percentage overlap against null spatial distribution at $P < 0.05$ after FDR correction across all networks.

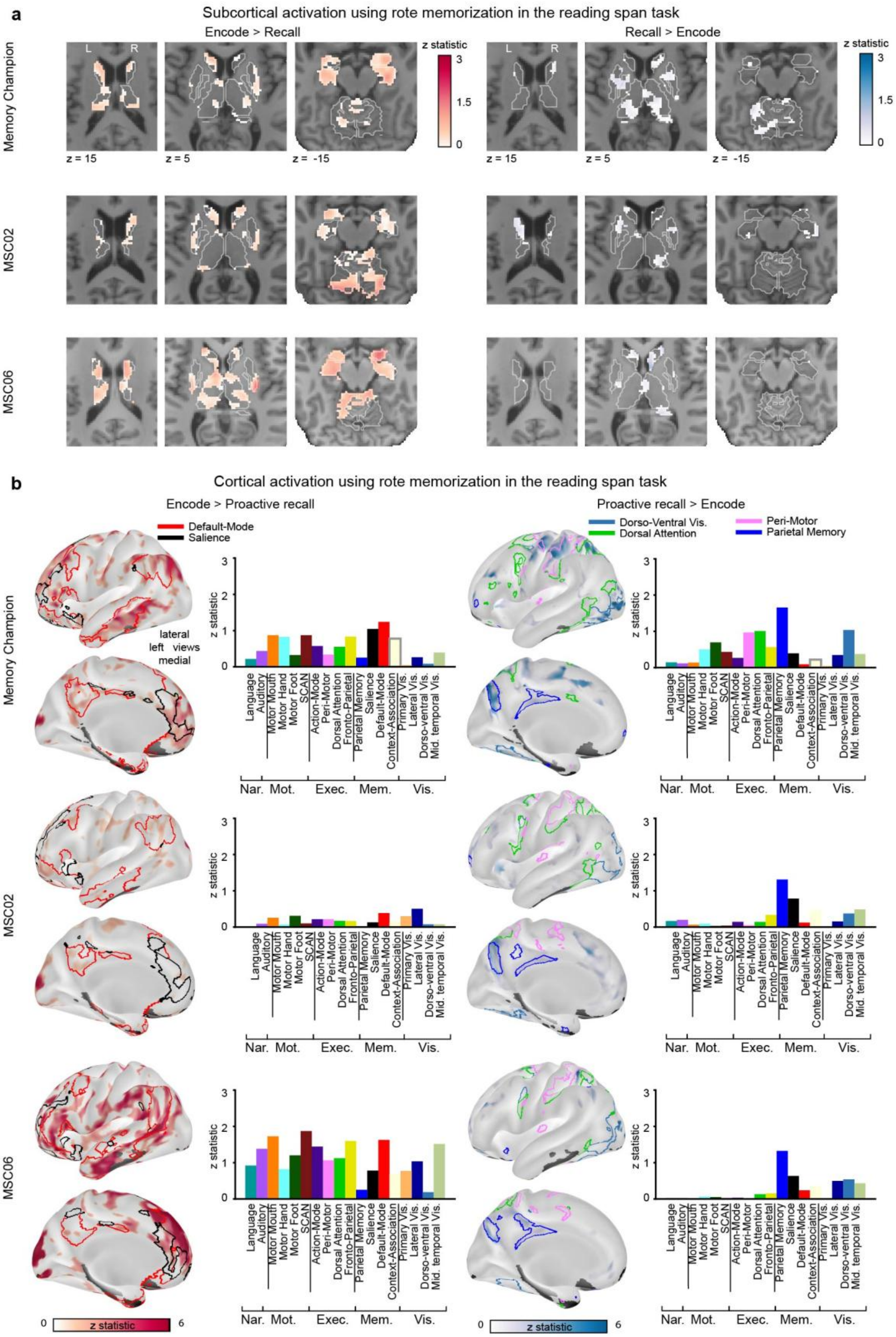
a
Subcortical activation using rote memorization in the reading span task
Encode > Recall
Recall > Encode
z statistic
3
1.5
0
L
R
z = 15
z = 5
z = -15
Memory Champion
MSC02
MSC06
b
Cortical activation using rote memorization in the reading span task
Encode > Proactive recall
Proactive recall > Encode
Default-Mode
Salience
Dorso-Ventral Vis.
Dorsal Attention
Peri-Motor
Parietal Memory
lateral
left views
medial
z statistic
Language
Auditory
Motor Mouth
Motor Hand
Motor Foot
SCAN
Action-Mode
Peri-Motor
Dorsal Attention
Fronto-Parietal
Parietal Memory
Salience
Default-Mode
Context-Association
Primary Vis.
Lateral Vis.
Dorso-ventral Vis.
Mid. temporal Vis.
Nar.
Mot.
Exec.
Mem.
Vis.
0
z statistic
6

**Extended Data Figure 8: Task fMRI activation in rote memorization in the Memory Champion and controls for Encode versus Recall contrasts. a)** Subcortical Encode versus Proactive Recall contrasts in the reading span task. Results are shown on the Memory Champion's, MSC02's, and MSC06's (top to bottom) T1 horizontal slices (z = 15, 5, -15) for unthresholded z-statistic maps of Encode > Proactive Recall (left) and Proactive Recall > Encode (right). White borders outline anatomical structures (Freesurfer segmentation). **b)** Top 20 percentile cortical task fMRI activation of Encode versus Proactive Recall contrast in the reading span task. Inflated left lateral and medial views of the top 20 percentile z-statistic cortical maps of the Memory Champion, MSC02, and MSC06 (top to bottom), as well as network average z-statistical values of the maps of Encode > Proactive Recall (left) and Proactive Recall > Encode (right). Network individualized borders significant in the Memory Champion are displayed on the inflated cortical views for comparison in controls: DMN and Salience on the left, and dorso-ventral visual, dorsal attention, peri-motor, and parieto-memory on the right. Abbreviations stand for: narrative (nat.), motor (mot.), executive (exec.), memory (mem.), visual (vis.), somato-cognitive-action (SCAN).

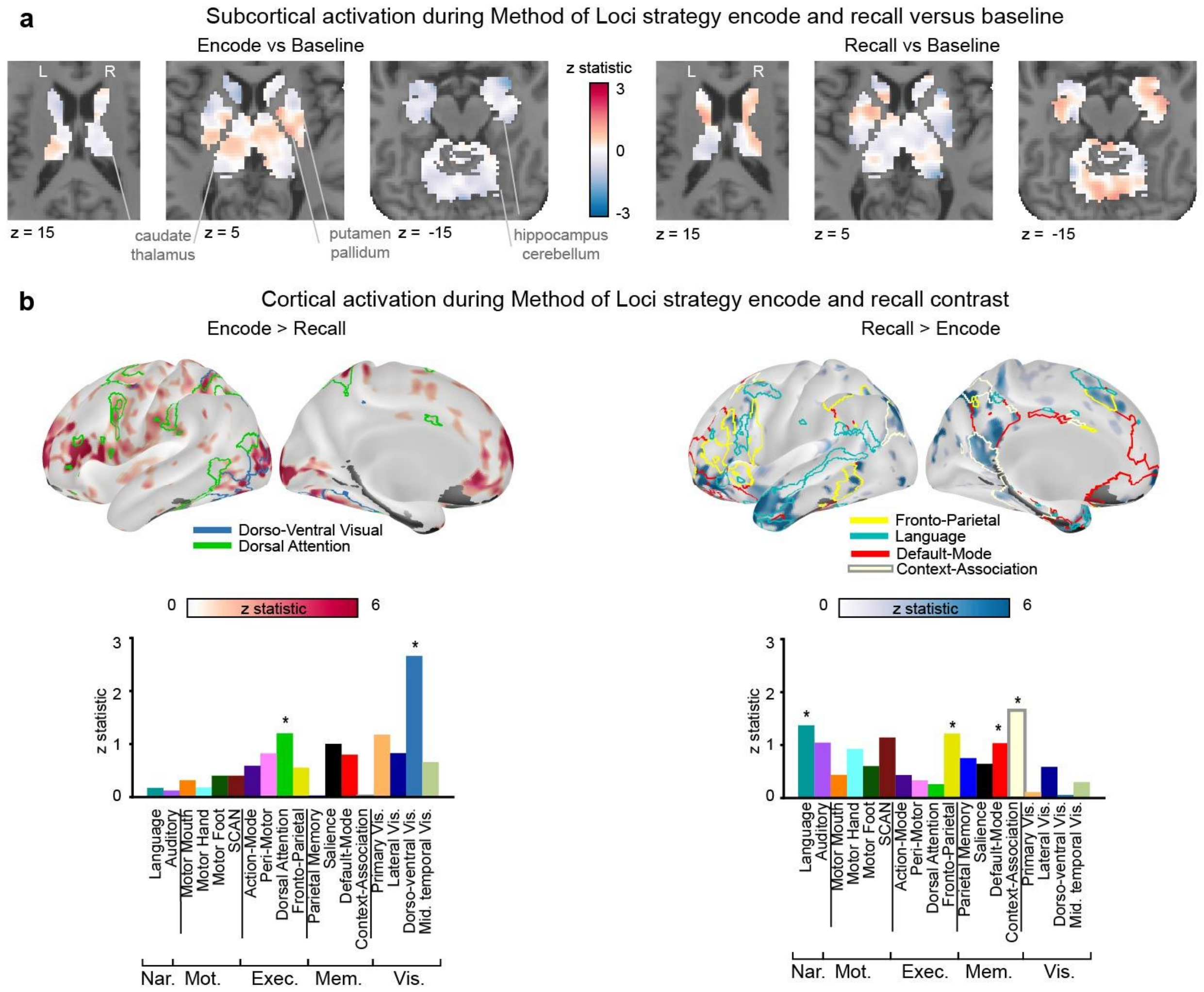


**Extended Data Figure 9: Difference in task fMRI activation between encoding and recall (proactive) using Method of Loci in the Memory Champion. a)** Task fMRI activation of Encode and Recall contrasts against baseline when the Memory Champion is using the Method of Loci strategy in the Deck of cards task. Results are shown on the Memory Champion's T1 horizontal slices (z = 15, 5, -15) for unthresholded Z-statistic maps of Encode > Baseline (left) and Recall > Baseline (right), only for subcortical anatomical structures. **b)** Top 20th percentile cortical activation of Encode versus Recall (Proactive) contrasts with the Method of Loci memory strategy in the deck of cards task for the Memory Champion. Inflated left lateral and medial views (top) and average per functional network (bottom) showing z-statistic cortical activation for Encode > Recall (left) and Recall > Encode (right). * indicates networks with significant z-statistics against the spatial null distribution at $P < 0.05$ after FDR correction across all networks. Significant network individualized borders are displayed on the inflated cortical views. Full cortical views are available in Supplementary Fig. 9. Abbreviations stand for: narrative (nat.), motor (mot.), executive (exec.), memory (mem.), visual (vis.), somato-cognitive-action (SCAN).

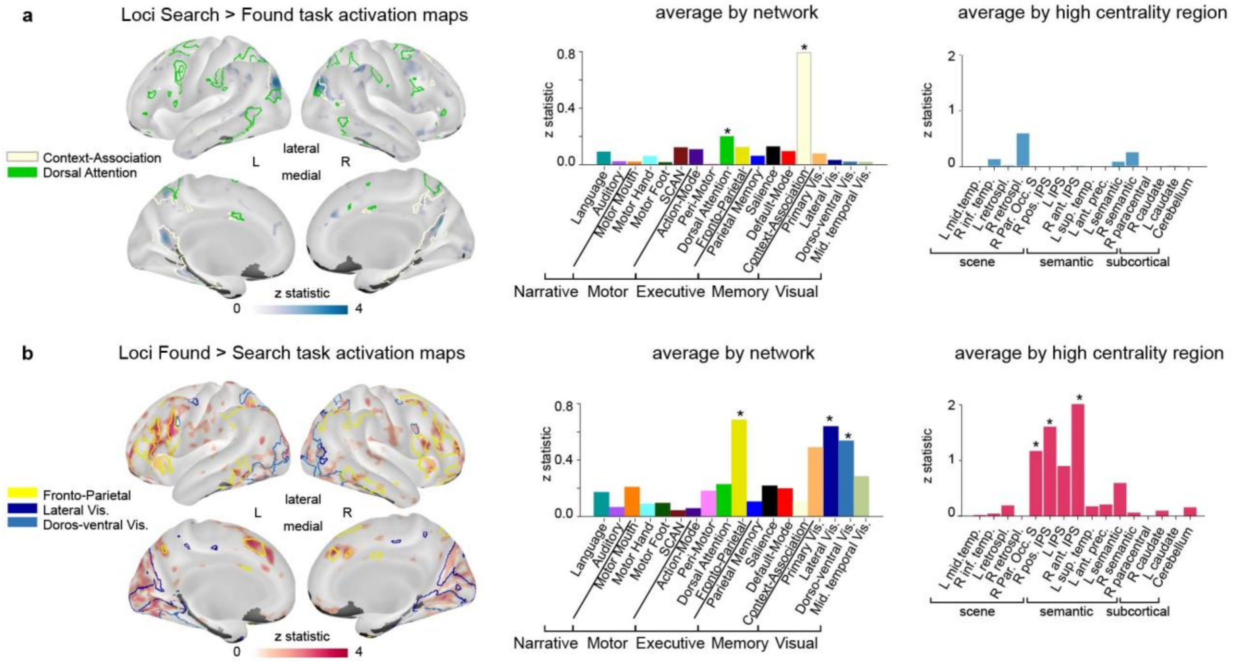


**Extended Data Figure 10: Task activation when recalling digits of the number pi using Method of Loci. a)** Memory Champion task activation contrast of Search > Found recall of 5-digit sequence locations within the 10,000 first digits of the number pi. Context-association and dorsal attention network borders are displayed on the inflated views. Average Z-statistic per network (middle) and per high centrality region (right) bar plot (*, $P < 0.05$, FDR-corrected). **b)** Memory Champion task activation contrast of Found > Search recall of 5-digit sequence locations within the 10,000 first digits of the number pi. Fronto-parietal, lateral visual and dorso-ventral visual network borders are displayed on the inflated views. Average Z-statistic per network (middle) and per high centrality region (right) bar plot (*, $P < 0.05$, FDR-corrected). Low signal-to-noise masked areas are displayed in grey.
