## Supplementary Figures for "Brain organization of a memory champion"

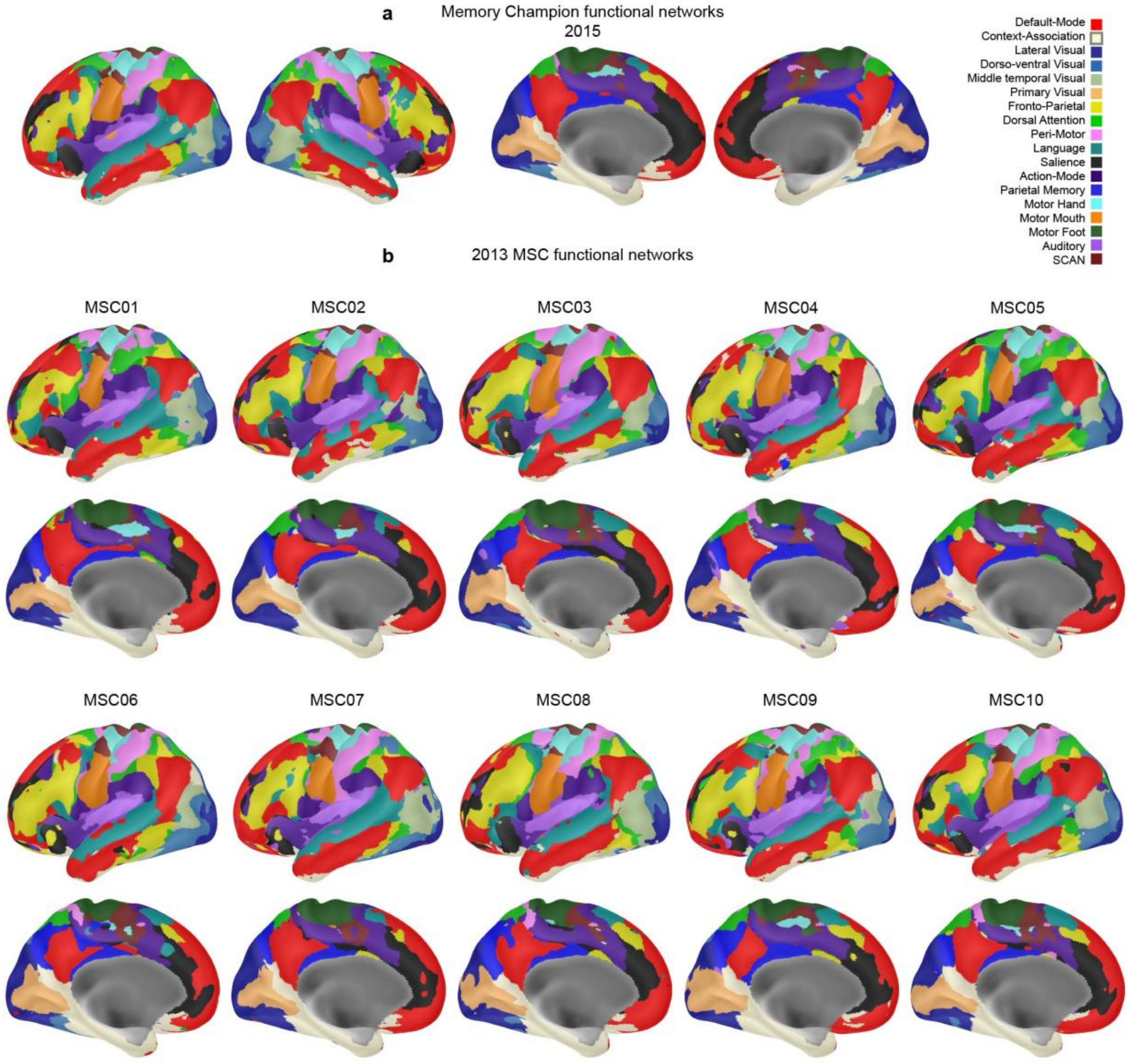


**Supplementary Figure 1: Individual-specific functional networks. a)** Full lateral and medial left and right views of the individual functional network from the 2015 resting state of the Memory Champion. **b)** Left lateral and medial inflated views of the individual network labeling for each of the MSC participants. Network color codes are displayed at the top right.

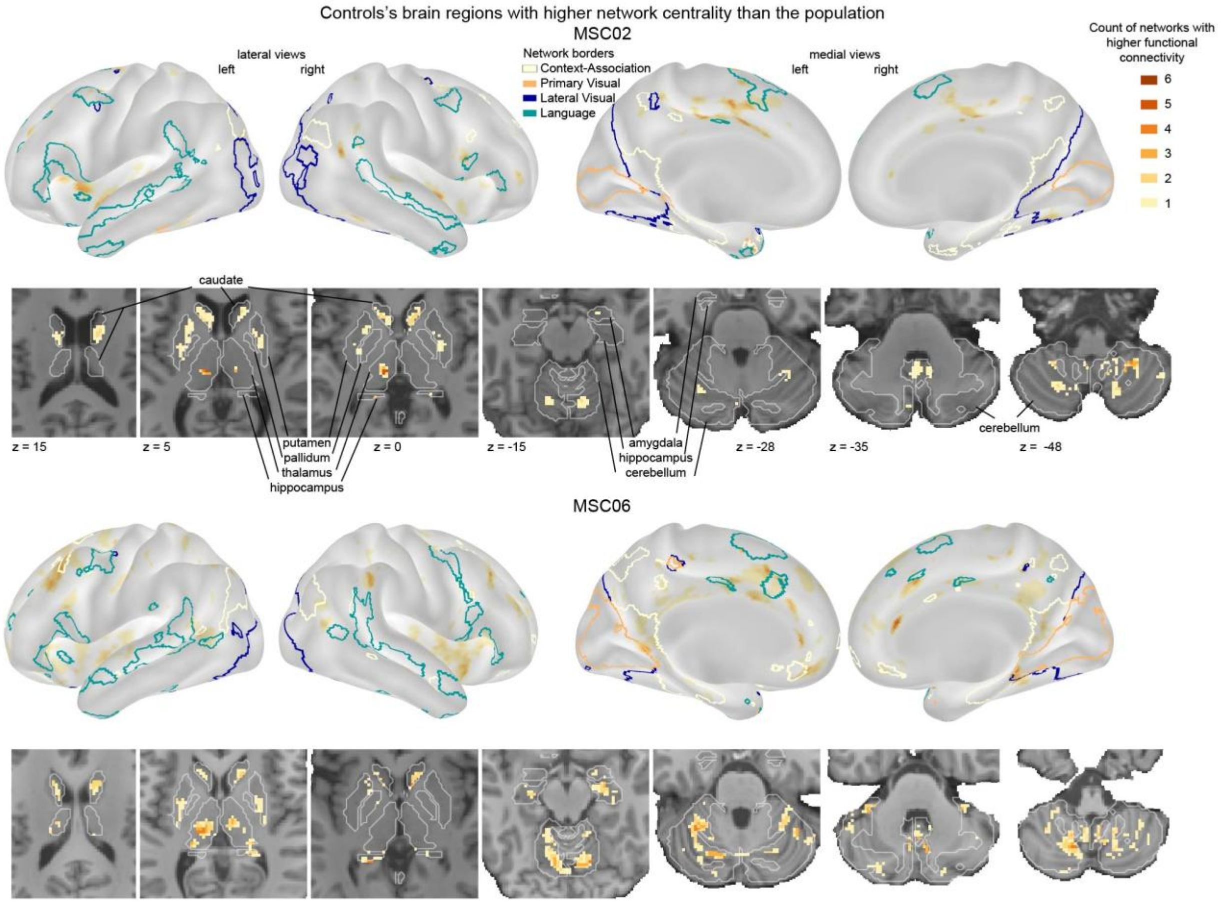


**Supplementary Figure 2: Brain regions with higher network centrality in MSC controls than the population**. Regions with higher functional connectivity in MSC02 (top) and MSC06 (bottom) than 95% of individuals from the HCP population, at both timepoints (MSC original acquisition and 2021), with one or more of the 17 functional networks after mixture modeling normalization of each network FC map (see Methods). Darker brown colors indicate stronger FC with a higher number of functional networks (from 1 to 6), indicating higher centrality. Boundaries for the language (teal), lateral visual (dark blue), primary visual (ochre), and context-association (CAN, white) networks are shown in color for comparison with the Memory Champion findings. Subcortical regions are displayed on the participants T1 horizontal slices and anatomical borders are displayed with white outlines.

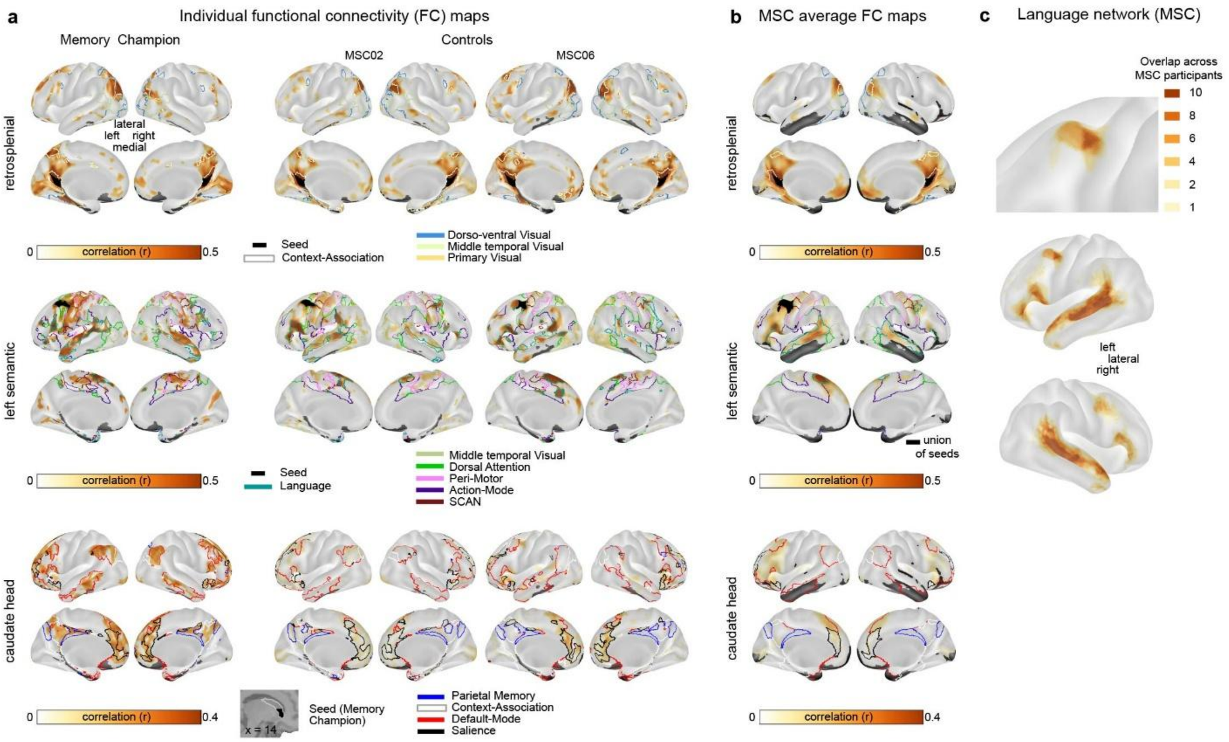


**Supplementary Figure 3: Full cortical views of the functional connectivity from high centrality regions. a)** Full cortical views of the whole-brain FC maps for representative brain regions identified in Fig. 1c, top 20% of strongest connections are shown. Left column shows the Memory Champion, middle column MSC02 control, and right column MSC06 control (2021 data; see Extended Data Fig. 2 for 2015 data, and Supplementary Fig. 3 for additional brain views). The top row shows correlation (r) for retrosplenial cortex (Brodmann areas 26, 29, and 30, part of context-association network -white border), middle row shows the left semantic region (area 55b; language network -teal border) and the bottom row shows the head of the left caudate. The seed regions are shown in black. All other network borders indicate networks significantly more connected in the Memory Champion compared to the controls in 2021. SCAN stands for somato-cognitive action network. Low signal-to-noise masked areas are displayed in grey. **b)** Full cortical views of the average whole-brain FC maps for representative brain regions identified in Fig. 1c for the MSC controls. The same regions as in panel A are displayed, showing the top 20% of the strongest connections. **c)** Overlap of the individual language network across MSC participants. The top panel is zoomed in on the left semantic region, while the middle and bottom panels show lateral views of the left and right hemispheres.

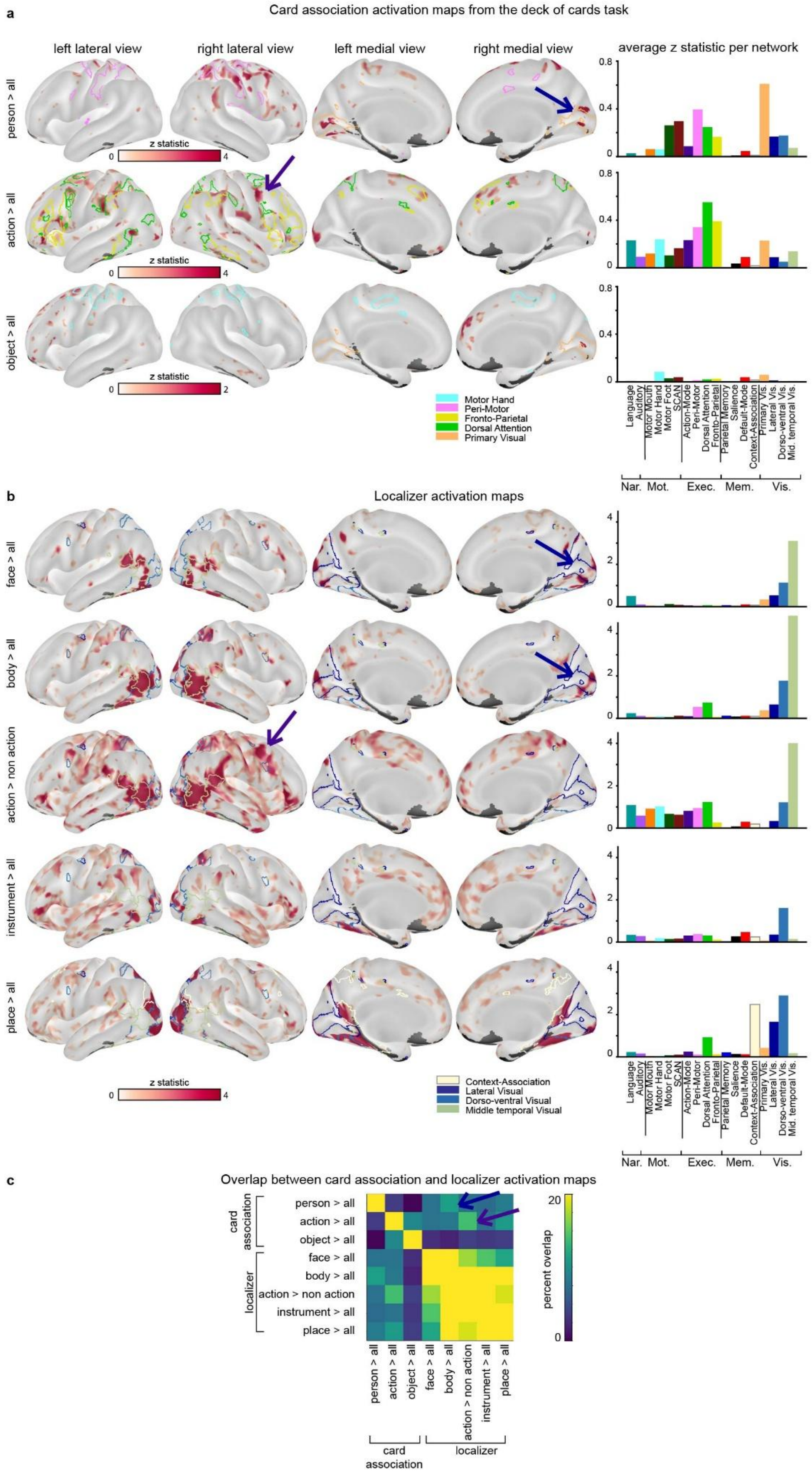

a
Card association activation maps from the deck of cards task
left lateral view
right lateral view
left medial view
right medial view
average z statistic per network
person > all
action > all
object > all
z statistic
0
4
2
0.8
0.4
0
Motor Hand
Peri-Motor
Fronto-Parietal
Dorsal Attention
Primary Visual
Language
Auditory
Motor Mouth
Motor Hand
Motor Foot
SCAN
Action-Mode
Peri-Motor
Dorsal Attention
Fronto-Parietal
Parietal Memory
Salience
Default-Mode
Context-Association
Primary Vis.
Lateral Vis.
Dorso-ventral Vis.
Mid. temporal Vis.
Nar.
Mot.
Exec.
Mem.
Vis.
b
Localizer activation maps
face > all
body > all
action > non action
instrument > all
place > all
Context-Association
Lateral Visual
Dorso-ventral Visual
Middle temporal Visual
c
Overlap between card association and localizer activation maps
card association
localizer
percent overlap
20
0

**Supplementary Figure 4: Pattern similarity in task fMRI activation maps of mental card association imagery and real item visualization in the Memory Champion. a)** Contrast between imagery of persons, actions and objects associated with cards, highlighting patterns specific to each item association in the Memory Champion. Card-associated person imagery particularly engages the visual primary network, while the dorsal attention network is particularly engaged in card-associated action imagery. **b)** Contrasts between the localizer tasks of each item against all others for face, body, instrument, and place, and for the independent localizer task of action, showing the action versus non-action trials contrast. (Left to right for panel a and b) left lateral, right lateral, left medial, and right medial inflated views of cortical activation contrast maps (Z-statistic) and average Z statistic value per functional network. The two network boundaries with the highest average activation values are displayed on the maps. Arrows point to activation area similarities between face, body localizers, and card association of person imagery, as well as action localizer and card-associated action imagery. **c)** Percent overlap between maps of panel a and b. Arrows highlight the most similar cortical patterns between card imagery and localizers.

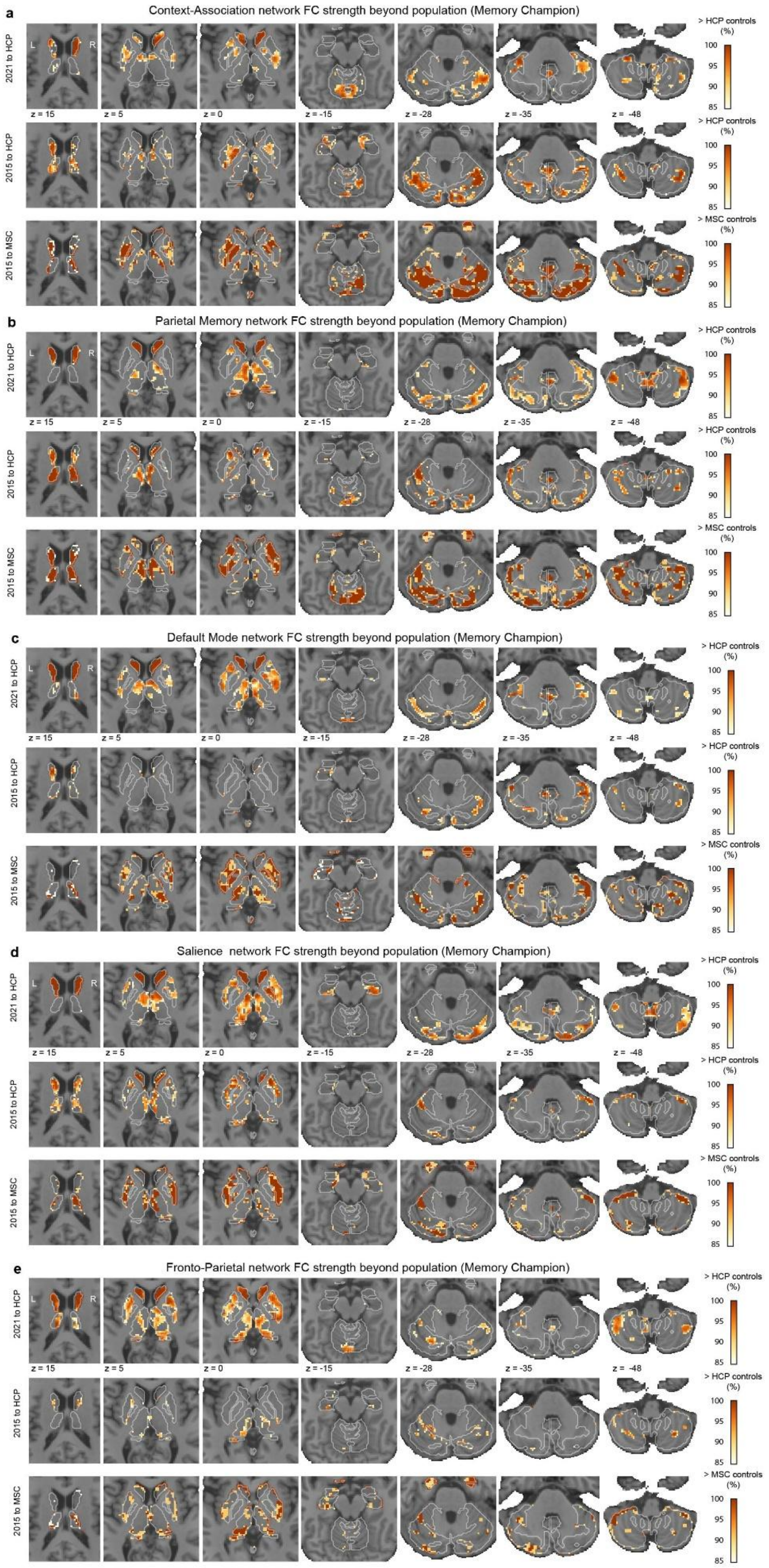
a
Context-Association network FC strength beyond population (Memory Champion)
b
Parietal Memory network FC strength beyond population (Memory Champion)
c
Default Mode network FC strength beyond population (Memory Champion)
d
Salience network FC strength beyond population (Memory Champion)
e
Fronto-Parietal network FC strength beyond population (Memory Champion)
2021 to HCP
2015 to HCP
2015 to MSC
L
R
z = 15
z = 5
z = 0
z = -15
z = -28
z = -35
z = -48
> HCP controls (%)
> MSC controls (%)
100
95
90
85

**Supplementary Figure 5: Stronger FC from subcortex to memory networks in the Memory Champion. a)** Subcortical regions where the Memory Champion showed stronger FC than controls with the Context-Association network. **b)** Subcortical regions where the Memory Champion showed stronger FC than controls with the Parietal Memory network. **c)** Subcortical regions where the Memory Champion showed stronger FC than controls with the Default-Mode network. **d)** Subcortical regions where the Memory Champion showed stronger FC than controls with the Salience network. **e)** Subcortical regions where the Memory Champion showed stronger FC than controls with the Fronto-Parietal network. For each panel, the first and second lines show percentiles beyond HCP controls for the 2021 and 2015 Memory Champion resting-state FC, respectively. The third line shows the FC percentile beyond MSC controls for the 2015 Memory Champion resting-state FC.

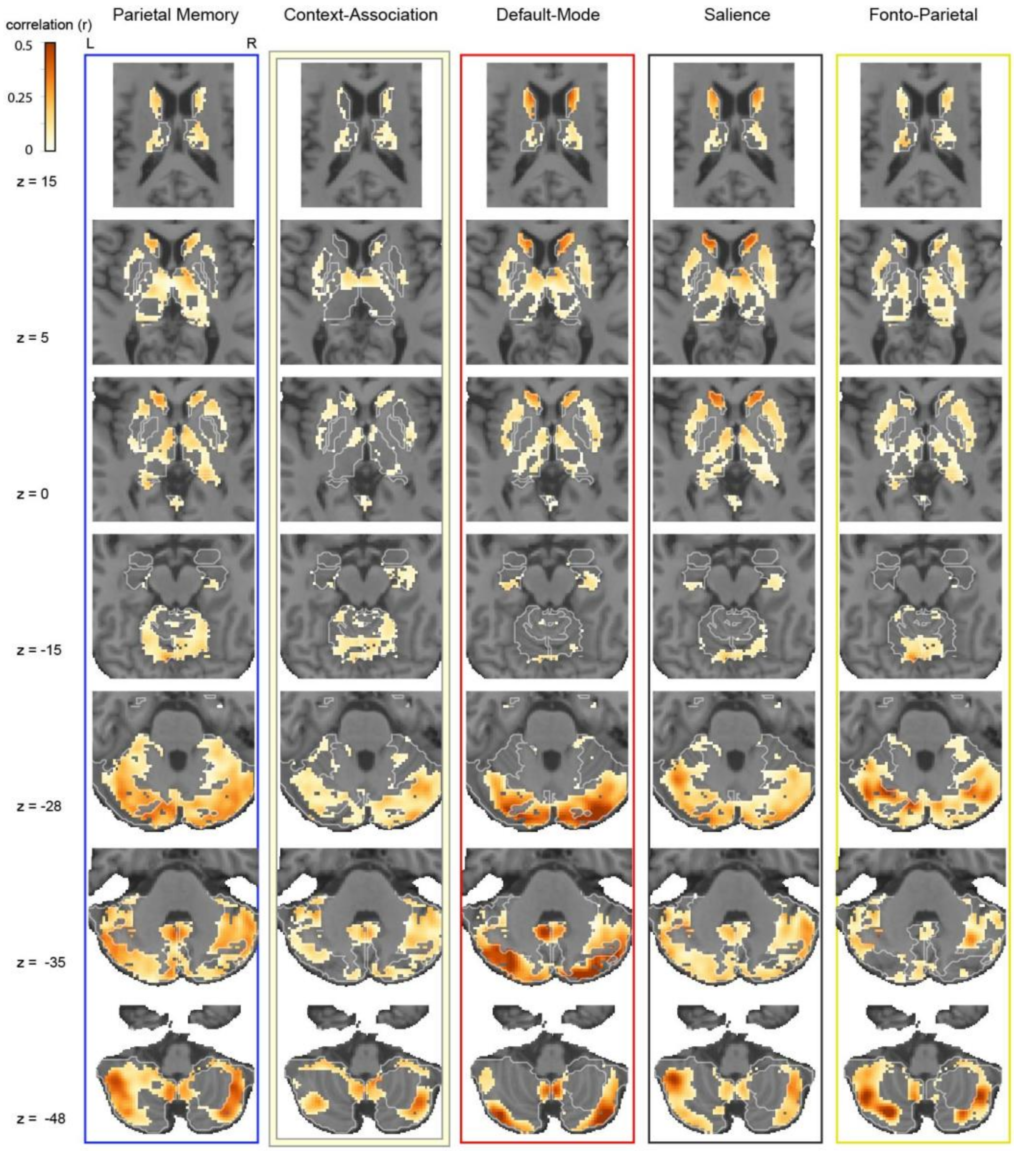


**Supplementary Figure 6: Subcortical network territory in the Memory Champion.** Horizontal slices show (left to right) the Parietal Memory, Context-Association, Default-Mode, Salience, and Fronto-Parietal network FC strength (correlation) for the subcortex in the Memory

Champion 2021 dataset. The network territory shows where a network is a competitor for winner-take-all labeling (see Methods). Results are displayed on the Memory Champion average T1.

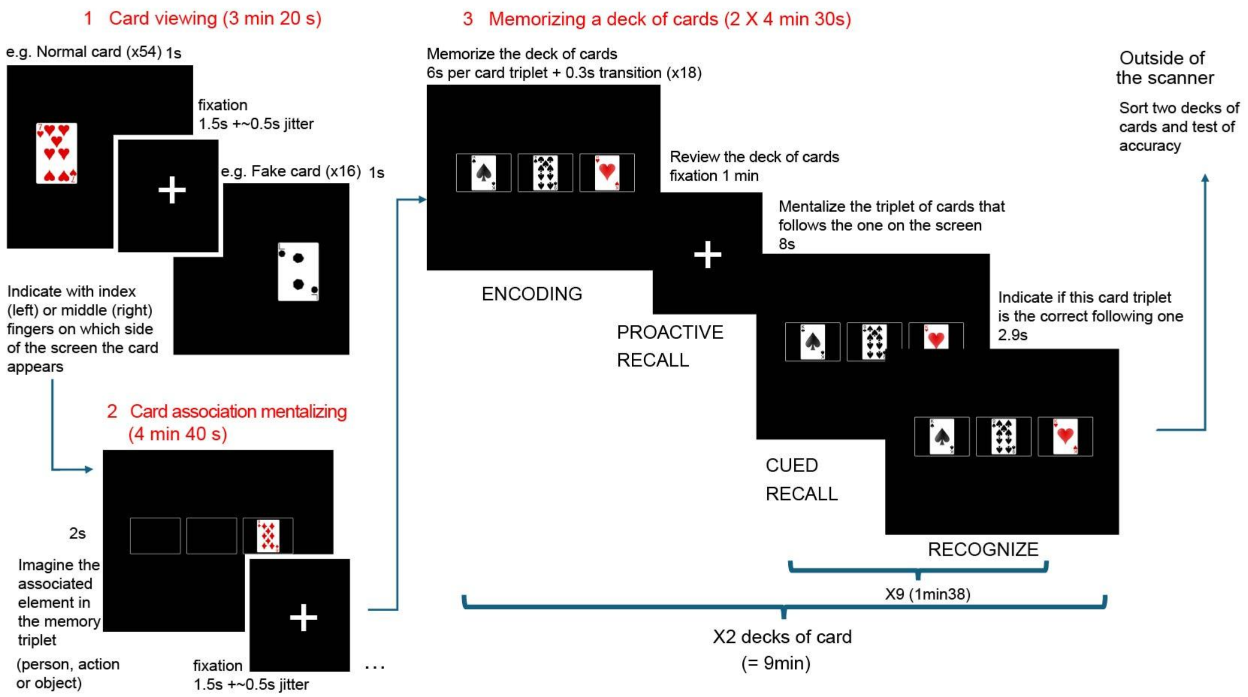


**Supplementary Figure 7: Deck of cards memory task schema.** The task is composed of three parts: 1) Visualizing a full deck of cards to obtain baseline card activation contrast. 2) Card association mentalizing. Mentalize the concept (person in position 1, action in position 2, object in position 3) associated with the card. Fake cards are also presented in each position. 3) Memorization and recall of a deck of cards. A deck of cards is presented in card triplets. A fixation cross appears for 1 minute during which the Memory Champion reviews the just-learned deck of cards order mentally. Then, 9 triplets are tested: a card triplet from the deck is shown, the Memory Champion must mentalize the card triplet following in the deck of cards, a new triplet appears, and the Memory Champion needs to judge if this is the correct following triplet of cards. This third part is repeated with another deck of cards. A second deck is memorized and tested. Then, the Memory Champion gets out of the scanner and reorders 2 physical decks of cards to verify the correct memorization.

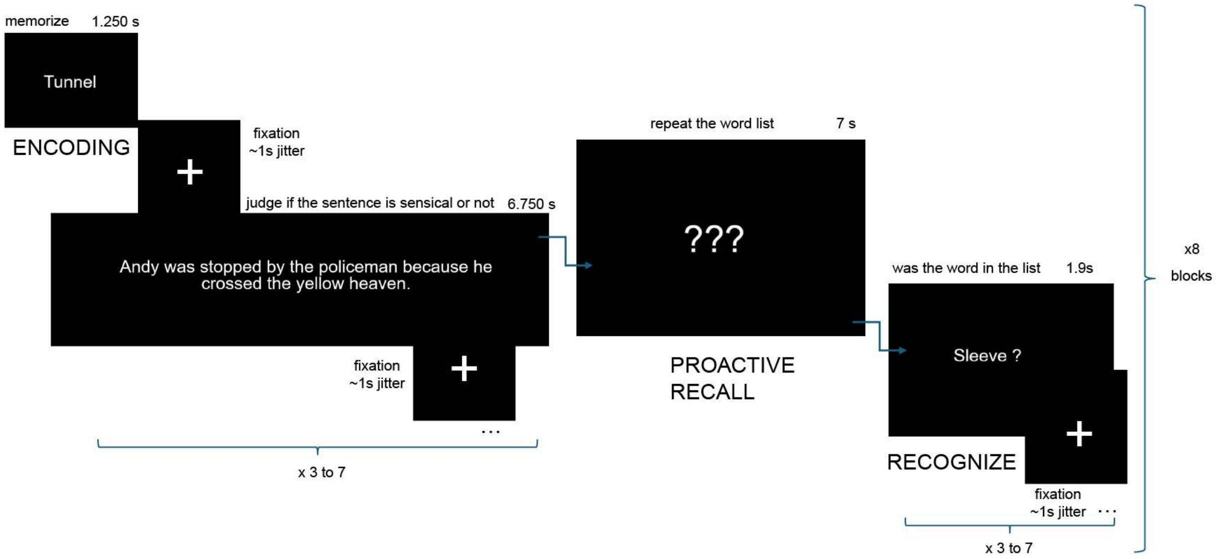


**Supplementary Figure 8: Reading span task schema.** This working memory task consists of the memorization of a list of 3 to 7 words. Words from the list to encode are alternated with sentences. The sentences need to be judged as sensical or not. After the list presentation, the participant is asked to mentally repeat the list of words. Then a list of words is presented, and the participant needs to answer if the word was in the list to memorize. Controls and the Memory Champion performed this task.

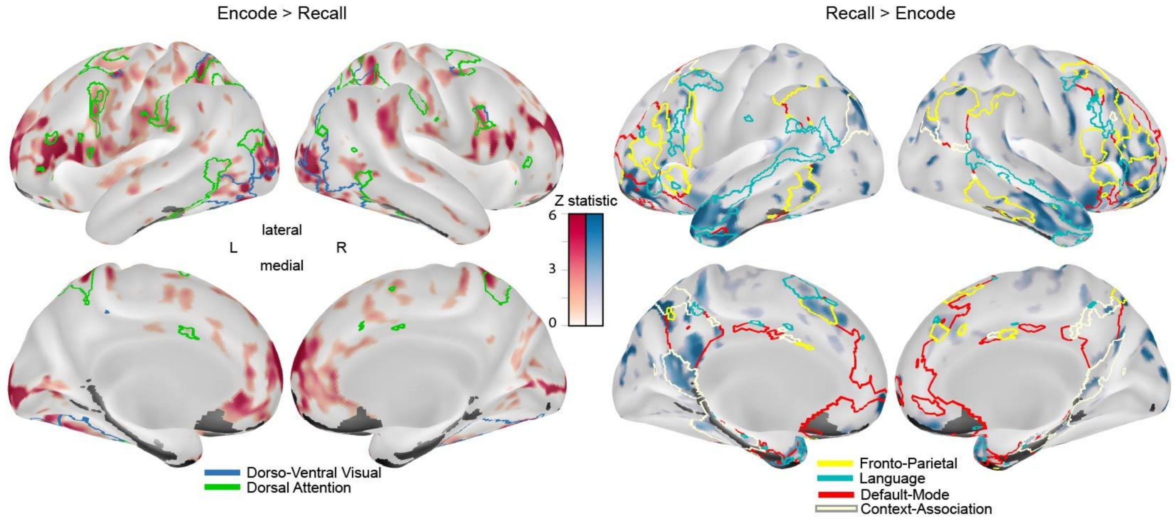


**Supplementary Figure 9: Difference in task fMRI activation between encoding and recall (proactive) using Method of Loci in the Memory Champion.** Top 20th percentile cortical activation of Encode versus Recall contrasts with the Method of Loci memory strategy in the deck of cards task for the Memory Champion. Inflated left lateral and medial views showing z-statistic cortical activation for Encode > Recall (left) and Recall > Encode (right). Significant network individualized borders are displayed on the inflated cortical views.

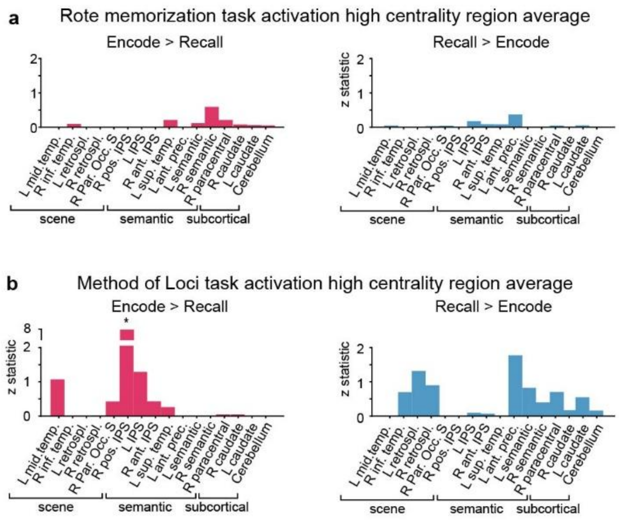


**Supplementary Figure 10: Rote memorization and Method of Loci encoding and recall contrasts in the Memory Champion**. **a)** Average Z statistics per high centrality regions in task fMRI contrast of Encoding versus Recall when using rote memorization in the Reading Span task. Positive z-statistic maps of Encode > Recall (left) and Recall > Encode (right). **b)** Average Z statistics per high centrality regions in task fMRI activation of Encode versus Recall contrasts when the Memory Champion is using the Method of Loci strategy in the Deck of Cards task. Positive z-statistic maps of Encode > Recall (left) and Recall > Encode (right) (*, one-tailed independent $t > 7.9$, $P < 0.001$, FDR). High centrality regions are grouped by scene and semantic modules and subcortical regions

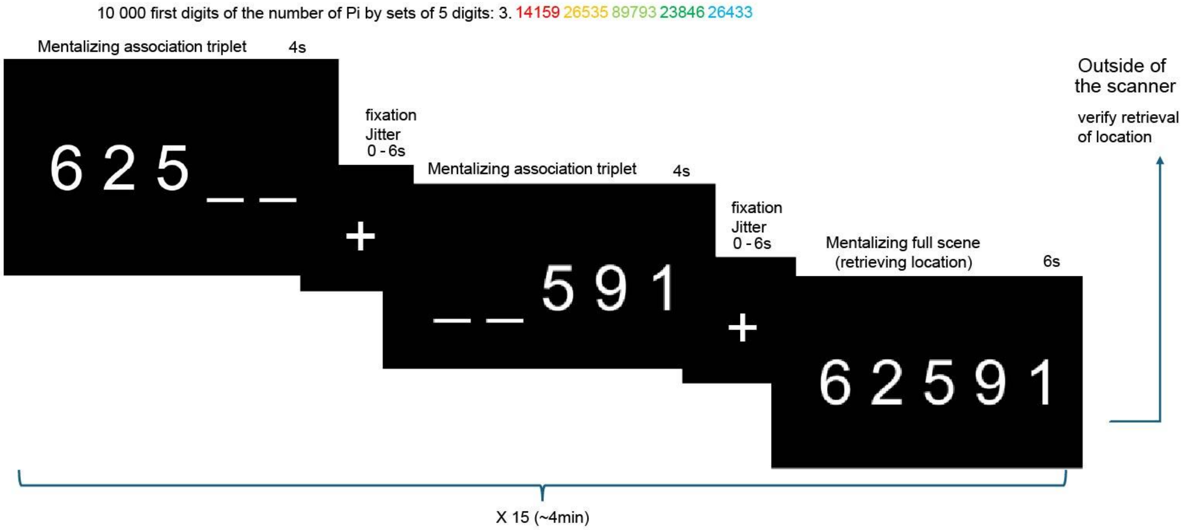


**Supplementary Figure 11: Number of pi task schema.** The Memory Champion had already memorized the first 10,000 digits of the number pi. The challenge he trained for was to recall the location of a sequence of 5 digits within the 10,000 first digits of pi and to retrieve the following 5 digits in the series. The memory method he used is based on paired association to digits of person, action and object (similar to cards for the speed card challenge), and using 2 triplets with one overlapping number to represent the sequence of 5 digits. We tested his mentalization of each triplet and his ability to retrieve the location of the full associated scene for 15 sequences of 5 digits. Outside of the scanner, the success or failure of retrieval of the location of each sequence was recorded.

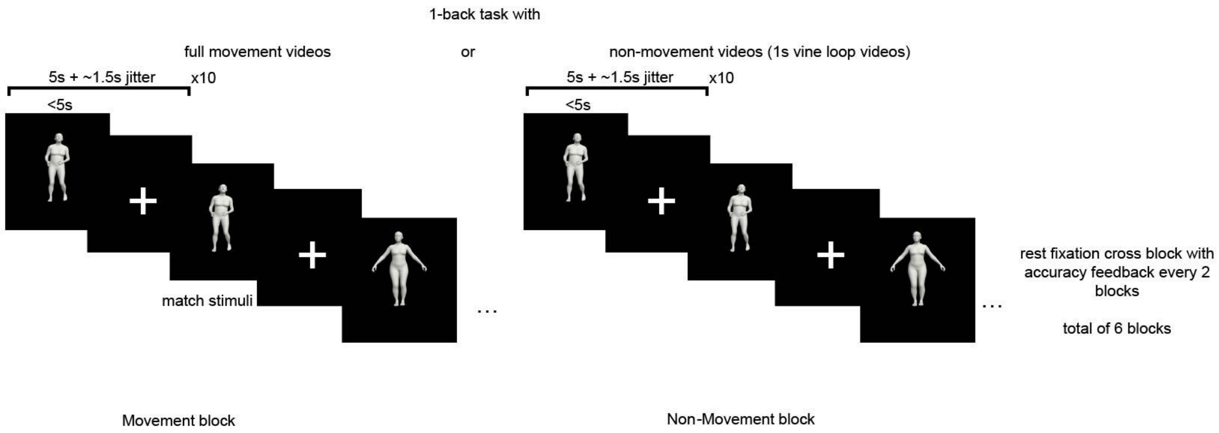


**Supplementary Figure 12: Action localizer task schema.** Movement videos from the MoVi (A Large Multipurpose Motion and Video, https://www.biomotionlab.ca/movi/) Dataset, representing 21 everyday actions and sport movements, are used in a 1-back block design. These are matched with the same database transformed into 1-second ping-pong loop videos (the first second back and forth repeated version) for non-movement match stimuli control.

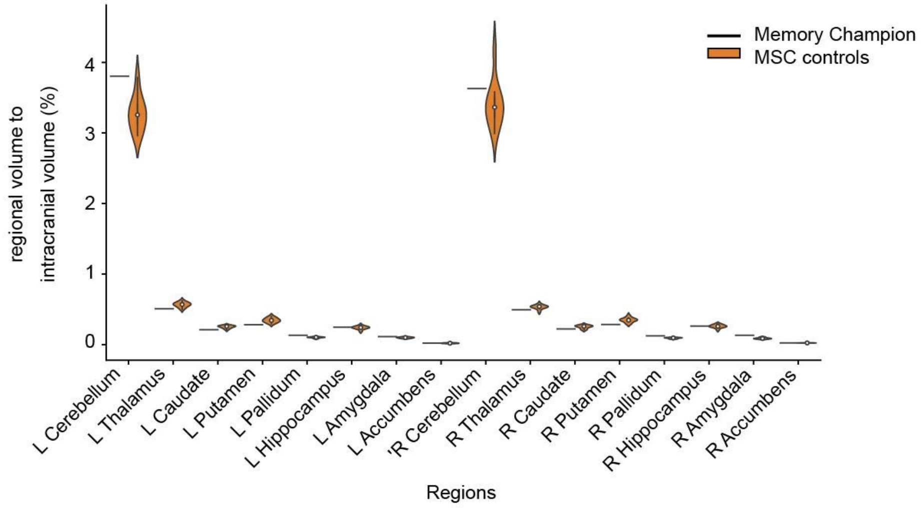


**Supplementary Figure 13: Regional structural volume of the Memory Champion and the MSC controls.** Distribution of the percentage regional volume compared to the intracranial volume of the cerebellum and subcortical regions segmented by Freesurfer in the left (L) and right (R) hemisphere.
